# Cranial suture integrity is maintained by Fgfr3 in zebrafish

**DOI:** 10.1101/2025.04.09.647929

**Authors:** Rachel Pereur, Yvan Marc, Yuliya Lim, Alain Schmitt, Marilyne Malbouyres, Anatole Chessel, Florence Ruggiero, Marie-Claire Schanne-Klein, Laurence Legeai-Mallet, Emilie Dambroise

**Affiliations:** Laboratory of Genetics of Developmental Disorders, Imagine Institute, Université de Paris Cité-INSERM UMR1163, Paris, France; Laboratory for Optics and Biosciences, Ecole polytechnique, CNRS, INSERM, Institut Polytechnique de Paris, Palaiseau, France; Université Paris Cité, CNRS, Inserm, Institut Cochin, F-75014 Paris, France; Laboratory of biology and pathology of the matrix, ENS of Lyon, Institut of functional genomics, UMR 5242, Lyon, France

## Abstract

Cranial suture formation is a dynamic process requiring precise cellular and molecular coordination to regulate bone growth and suture homeostasis. Despite the recognized role of Fibroblast Growth Factor Receptor 3 (FGFR3) in this process and its association with Muenke syndrome, one of the most common syndromic craniosynostoses, its precise function remains undefined. Through analyses of cranial suture formation and maintenance in a *fgfr3* LoF zebrafish model displaying abnormal suture morphology, we demonstrate for the first time that Fgfr3 plays a pleiotropic role during these events. Transmission electron microscopy and second harmonic generation imaging revealed that Fgfr3 is involved in the proper organization of the collagen network within the suture. By employing specific transgenic reporter lines and studying gene expression of suture mesenchymal stem cell markers via RNAscope in situ hybridization, we show that Fgfr3 is crucial for regulating osteogenesis within the suture. Specifically, Fgfr3 limits the number of osteoprogenitors at the osteogenic front and promotes osteoblast maturation at the suture edge. Finally, our results reveal that Fgfr3-mediated regulation of osteogenesis involves cross-talk between FGF, canonical Wnt, and potentially BMP signaling pathways. In conclusion, these data position Fgfr3 as a central regulator of cranial suture formation and open new perspectives for understanding suture homeostasis and FGFR3-related craniosynostoses.

## Introduction

Cranial sutures are essential structures in skull development. Acting as fibrous joints between rigid calvarial bones, they provide plasticity for skull expansion during early brain growth and contribute to skull shape, symmetry, and absorption of mechanical forces through adulthood ^1^. In humans, mice, and zebrafish, five primary sutures extend from anterior to posterior, delineating the boundaries between cranial bones: the metopic suture, the two coronal sutures, the sagittal suture, and the lambdoid suture ^2^. Their formation is a dynamic process requiring precise cellular and molecular coordination to regulate bone growth and suture homeostasis. At the edges of the approaching bones, an osteogenic front (OF) composed of osteoprogenitor cells is established. These proliferating cells sequentially differentiate into immature and mature osteoblasts forming new bone matrix that is initially restricted to growing bone tips and then at the edges of the suture ^3–5^. To support this process, a heterogeneous population of suture mesenchymal stem cells (SuSCs), expressing various markers such as Gli1, Axin2, Prrx1, Ctsk, Bmpr1a, Ddr2, and Grem1a, resides between the OFs, preserving suture patency in a fibrous, unossified state while also contributing to the recruitment of osteoprogenitors at the ossification front. Each subpopulation presents their own properties and participates more or less in the osteogenesis within the OF ^5–12^. All these cells are surrounded by an extracellular matrix (ECM) rich in collagens, which provides structural support and participates in the transmission of biomechanical signals, as sutures are subjected to significant tensile and compressive forces, both essential for the regulation of osteogenesis ^13–18^.

Several signaling pathways and genes involved in suture homeostasis have been implicated in genetic diseases characterized by pathological suture expansion or premature fusion, known as craniosynostosis. This includes Fibroblast Growth Factor Receptor 3 (FGFR3), where both gain-of-function (GoF) and loss-of-function (LoF) mutations lead to distinct cranial suture defects. GoF mutations are associated with syndromic craniosynostosis, as observed in Muenke syndrome, one of the most frequent forms of craniosynostosis, and in Crouzon syndrome with acanthosis nigricans^19,20^. In contrast, LoF mutations result in Camptodactyly-Tall Stature-Scoliosis-Hearing Loss (CATSHL) syndrome, which leads to the formation of ectopic bones along the sutures, known as Wormian bones ^21^. However, despite these findings, *Fgfr3* mouse models with GoF or LoF mutations have failed to fully recapitulate these cranial phenotypes, posing a significant challenge in deciphering the precise role of FGFR3 in cranial suture formation and maintenance ^22–24^. Zebrafish, now widely used to study craniofacial development in normal and pathological contexts (including craniosynostoses), has emerged as a relevant model for investigating Fgf signaling and Fgfr3 function during cranial vault formation ^5,11,25,26^. Similar to CATSHL syndrome, *fgfr3* LoF zebrafish exhibit microcephaly and Wormian bones ^27,28^. Our previous data on a *fgfr3* LoF zebrafish model (*fgfr3^lof/lof^)* demonstrated that *fgfr3* is expressed during cranial bone development, in late-stage osteoprogenitors and both immature and mature osteoblasts, and regulates positively both the expansion and maturation of immature osteoblasts ^28^. Interestingly, *fgfr3^lof/lof^* zebrafish exhibit a dramatic impairment of metopic and sagittal sutures formation, with bones that fail to overlap and remain separated by a thick layer of fibrous tissue. This phenotype, which contrasts with craniosynostosis, makes *fgfr3^lof/lof^* fish an ideal model for studying the involvement of Fgfr3 in cranial suture homeostasis.

Here, to achieve this goal, we analyzed the key factors involved in the formation and maintenance of cranial sutures in *fgfr3^lof/lof^* zebrafish, from suture initiation at 12 standard length (SL) to adulthood. We show that the absence of Fgfr3 completely disrupts suture formation, affecting not only the structural integrity of the sutures but also the organization of the collagen network within the sutures and at the bone ends, as shown by transmission electron microscopy (TEM) and second harmonic generation (SHG) microscopy. In addition, using transgenic lines marking osteoprogenitors (Tg: (*runx2;*GFP)), and immature (Tg:(*sp7;*mCherry)) and mature (Tg: (*bglap;*GFP)) osteoblasts, we demonstrate that the absence of Fgfr3 affects the entire membranous ossification process involved in suture formation and maintenance. Finally, by RNAscope *in situ* hybridization of SuSC marker expression, we provide evidence that the absence of Fgfr3 influences osteogenesis through the activation of the canonical Wnt pathway and potentially the BMP pathway.

## Results

### Fgfr3 is required for proper cranial suture formation

In adult wild-type zebrafish, metopic, sagittal and coronal sutures (Figure 1A) present the same structure as human coronal sutures, with overlapping bones, osteogenic cells at their edge and a thin layer of SuSCs surrounded by ECM rich in collagens in the middle. We previously reported that *fgfr3^lof/lof^* metopic and sagittal sutures displayed a failure of bone overlap, with the bones being separated by a thick fibrous tissue at 20 SL ^28^. As the metopic suture is among the first to develop in zebrafish, we focused on the role of Fgfr3 during formation of this suture. We used RNAscope *in situ* hybridization to map *fgfr3* expression during metopic suture formation in wild-type fish at three stages: 10 SL (frontal bones approaching), 12 SL (suture formation), and 20 SL (suture maintenance, adult stage) (Figure 1B, C). At 10 SL, *fgfr3* was expressed at the OF of the developing cranial bones, and on the forming metopic suture (Figure 1B). At later stages, transversal sections showed *fgfr3* expression within the suture and in osteogenic cells above and below the frontal bones, a region we will refer to as the bone surface (Figure 1C). At 12 SL, we noted that *fgfr3* expression was higher within the suture compared to the bone surface, while at 20 SL, the same level of *fgfr3* expression is observed within the suture and along the bone surface (Figure 1D).

**Figure 1.**
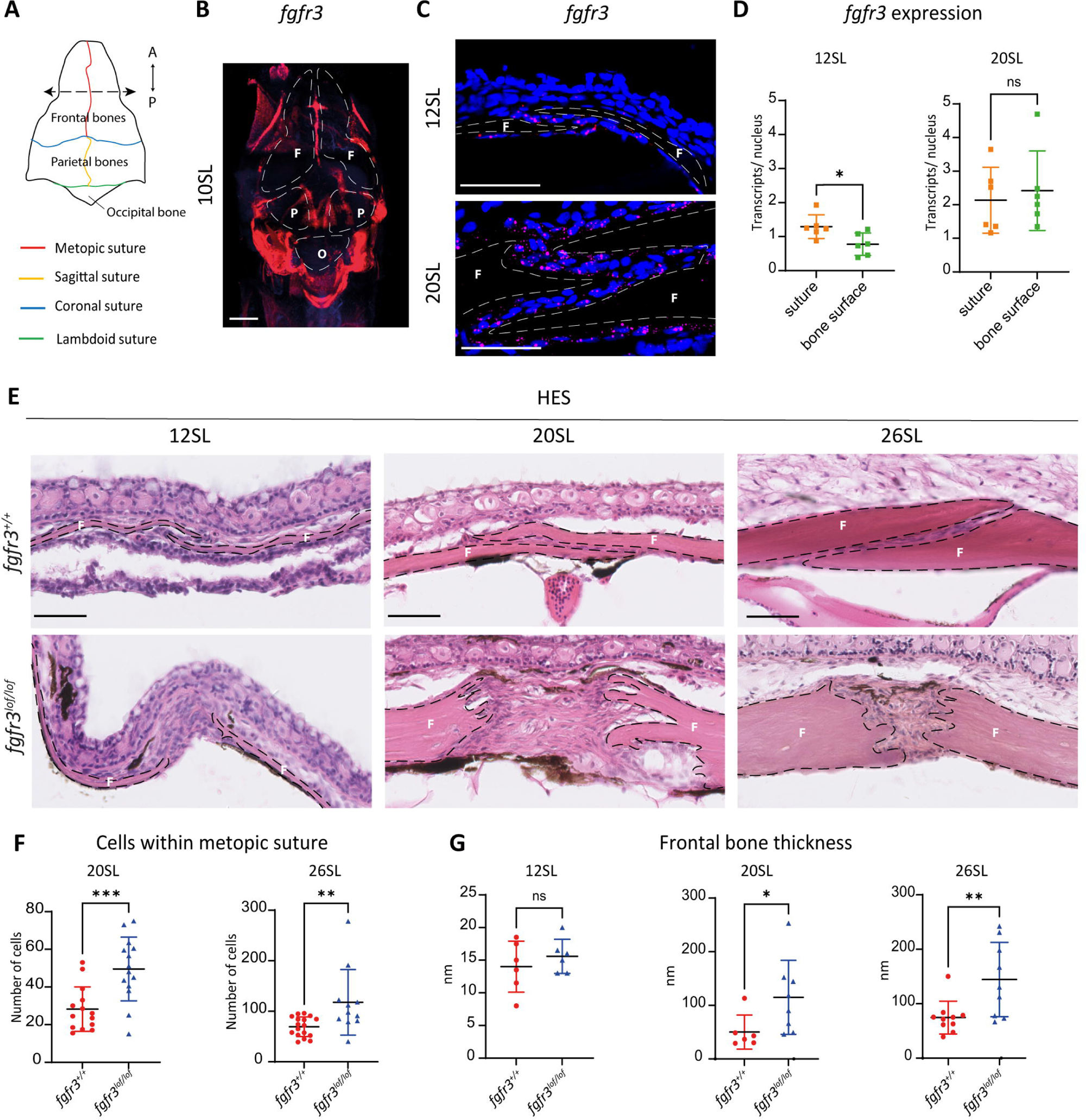
Fgfr3 is required for cranial vault suture formation. (A) Schematic representation of the zebrafish cranial vault. The dotted arrow corresponds to the plane of sectioning. (B) *fgfr3* expression (red) on whole mount *fgfr3^+/+^* cranial vault at 10 SL labelled by RNAscope *in situ* hybridization showed the high expression of *fgfr3* at the periphery of cranial bones and in the forming sutures. F= Frontal bone, P= Parietal bone, O= Occipital bone. Scale bar = 300 µm. (C) *fgfr3* expression (red) on coronal sections of *fgfr3^+/+^* metopic suture at 12 SL and 20 SL labelled by RNAscope *in situ* hybridization. At both stages, *fgfr3* was expressed in cells along the bone surface and within the suture. Nuclei were counterstained with DAPI (blue). Scale bar = 50 µm. (D) RNAscope, quantitative analyses of the number of transcript (dot) relative to the number of nuclei (DAPI) highlighted that at 12 SL *fgfr3* was more highly expressed inside the suture compared to the bone surface. In contrast, at 20 SL, no difference of *fgfr3* expression was observed based on localisation (12 SL and 20 SL: *fgfr3^+/+^* n=6; *fgfr3^lof/lof^* n=6). (E) HE stained coronal sections of the zebrafish head at the metopic suture level at 12, 20 and 26 SL demonstrated that the absence of Fgfr3 induced a delay of frontal bone growth, and prevented frontal bones overlap at 20 and 26 SL. The dotted lines represent the bone boundaries. Scale bar = 50 µm. (F) Quantification of the number of cells inside the suture at 20 SL (*fgfr3^+/+^* n=14; *fgfr3^lof/lof^* n=14) and 26 SL (*fgfr3^+/+^* n=17; *fgfr3^lof/lof^* n=11) showed that the *fgfr3^lof/lof^* fish present a greater number of cells within the suture at adulthood. (G) Measurement of frontal bone thickness close to the metopic suture at 12 SL (*fgfr3^+/+^* n=6; *fgfr3^lof/lof^* n=6), 20 SL (*fgfr3^+/+^* n=6; *fgfr3^lof/lof^* n=8) and 26 SL (*fgfr3^+/+^* n=10; *fgfr3^lof/lof^* n=9) showed that the *fgfr3^lof/lof^* bones are thicker than *fgfr3^+/+^* bones at adulthood. F: frontal bone. Data are presented as mean ± SD. The p-values were determined by Student’s t tests: ns: not significant; p < 0.05 = *; p < 0.01 = **; p < 0.001 = ***.

To further investigate the progression of the metopic suture in *fgfr3^lof/lof^* fish over time, we examined it at 12, 20 and 26 SL (Figure 1E). While *fgfr3^+/+^* fish displayed a well-formed suture at 12 SL, *fgfr3^lof/lof^* frontal bones remained separated in agreement with our previous data ^28^. By 20 and 26 SL, although the bones sometimes came closer together, they still did not overlap in the mutants (Figure 1E). Interestingly, lack of bone overlap was associated with a greater cell number within the *fgfr3^lof/lof^* metopic suture compared to controls (20 SL: *fgfr3^+/+^*: 28.25 ± 11.82; *fgfr3^lof/lof^*: 49.57 ± 16.95 p=0.0007 and 26 SL: *fgfr3^+/+^*: 69.41 ± 19.52; *fgfr3^lof/lof^*: 117.77 ± 64.94 p=0.0060) as well as a greater thickness of *fgfr3^lof/lof^* bones compared with *fgfr3^+/+^* bones in adulthood (+129% at 20 SL and +118% at 26 SL in the *fgfr3^lof/lof^* fish vs *fgfr3^+/+^* fish) (Figure 1F-G).

Interestingly, by performing sagittal sections of *fgfr3^lof/lof^* heads at 20 SL, we observed that coronal sutures are also affected, suggesting that the absence of Fgfr3 impacts all cranial vault sutures in zebrafish similarly (Figure 1 supplement 1A). Increased bone thickness in *fgfr3^lof/lof^* zebrafish was also observed in the bones forming the coronal suture (+308% at 20 SL and +164% at 26 SL) (Figure 1 supplement 1B).

These findings demonstrate that the absence of *fgfr3* disrupts proper cranial suture formation throughout the lifespan of zebrafish and leads to bone expansion, predominantly in thickness rather than length.

### The absence of Fgfr3 leads to a progressive disruption of Collagen fibrils within the suture

Cells within cranial vault sutures of *fgfr3^lof/lof^* fish are surrounded by a dense ECM rich in collagens, as shown by Sirius red-Alcian blue staining (Figure 2A). In our previous study, we found that the absence of Fgfr3 during cranial vault development leads to the upregulation of the expression of several ECM-regulating genes, including *col6a1*, *col12a1a*, and *col12a1b* ^28^. To determine whether this overexpression persisted in sutures, we performed immunofluorescence for ColVI,ColXII, and ColI which is also known to be expressed in cranial sutures. We observed that ColI and ColXII were expressed within the cranial sutures of both *fgfr3^+/+^* and *fgfr3^lof/lof^* fish at 12 SL, and that the expression levels of both proteins are increased only in older *fgfr3^lof/lof^* fish (20 SL) (Figure 2B-C). In contrast, ColVI was expressed to a lesser extent at both stages and genotypes (Figure 2D). As ColXII is known to interact with ColI fibrils and to regulate collagen fibril spacing and assembly during fibrillogenesis, we further analyzed collagen fibrils within the metopic suture by TEM at 9, 12 and 20 SL (Figure 2E) ^29^. At 9 SL, metopic sutures were not formed, and collagen fibrils were poorly organized for both genotypes. By 12 SL, metopic sutures in controls were formed with organized fibril clusters, while in mutants, where frontal bones remained separated, fibrils remained disorganized, similar to 9 SL. At 20 SL, fibrils in *fgfr3^lof/lof^* fish were clustered, similar to controls, but appeared less organized, with the size of each fibril appearing more heterogeneous (Figure 2E). This observation was confirmed by measurement of the transverse area of each fibril (Figure 2F). In controls, fibril size progressively increased over time, with most fibril areas not exceeding 1000 nm² at 20 SL. In *fgfr3^lof/lof^* zebrafish, we first observed no change in fibril size between 9 and 12 SL, consistent with the delay in frontal bone closure. Then, at 20 SL, significant heterogeneity in fibril size was observed, with larger fibrils emerging, exceeding 2500 nm². This increase was further confirmed by the larger average collagen fibril area for *fgfr3^lof/lof^* at 20 SL compared to controls (*fgfr3^+/+^*: 794.5 ±123.1 nm²; *fgfr3^lof/lof^*: 945.1 ±146.1nm²; p=0.05), showing an impairment of fibrillogenesis in adulthood within the suture in the mutants (Figure 2G).

**Figure 2.**
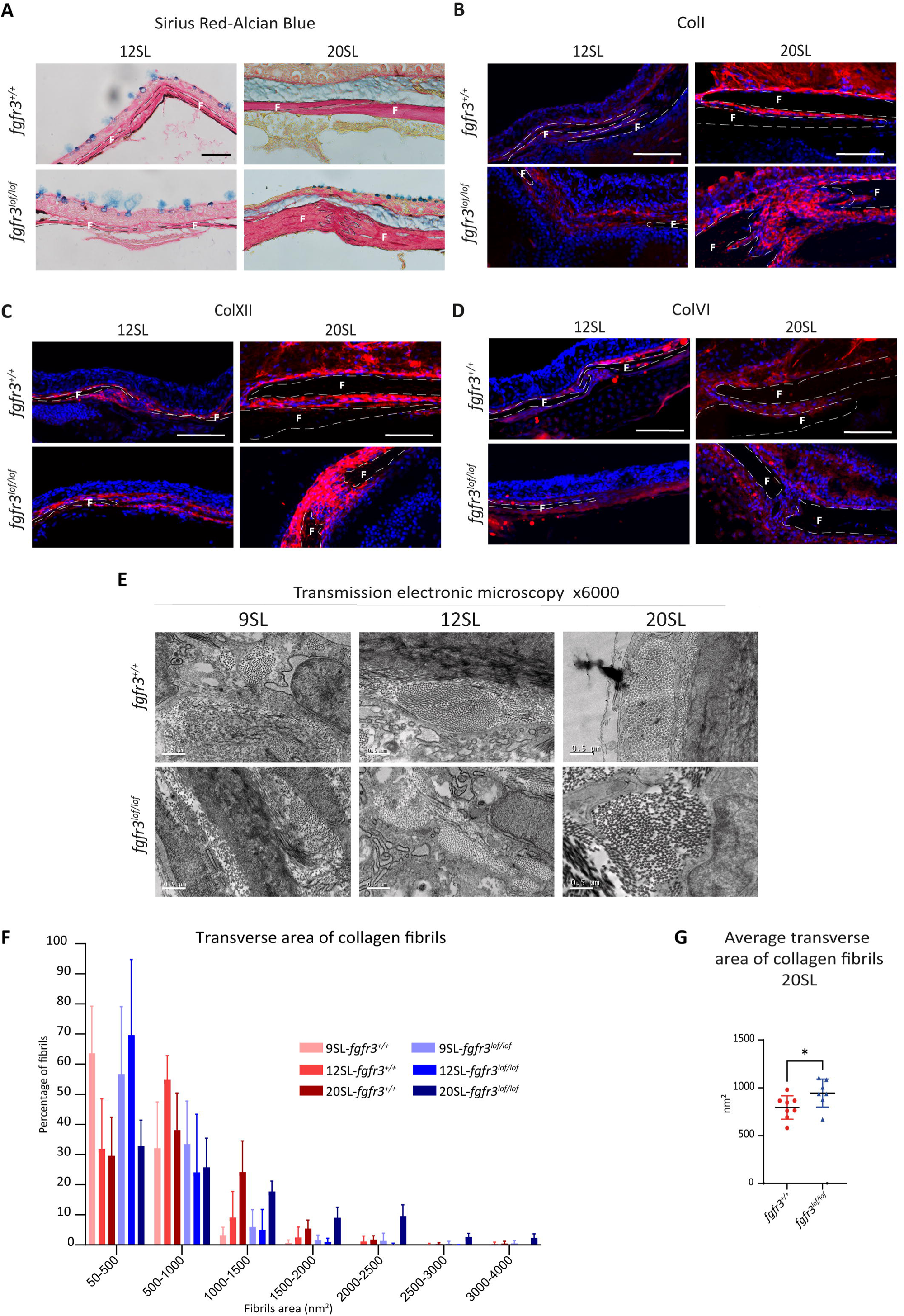
Fgfr3 deficiency leads to disruption of fibrillogenesis within the suture at adulthood. (A) Sirius Red–Alcian Blue stained coronal sections of the zebrafish head at the metopic suture level at 12 and 20 SL demonstrated that the absence of Fgfr3 induced an accumulation and a disorganization of the suture ECM at 20 SL. The dotted lines represent the bone boundaries. Scale bar = 50 µm. (B-D) Immunofluorescence against ColI (B), ColXII (C) and ColVI (D) on metopic sutures of *fgfr3^+/+ or lof/lof^* fish at 12 and 20 SL showed a higher amount of ColI and ColXII within the suture of the *fgfr3^lof/lof^* fish. Nuclei were counterstained with DAPI (blue) and the dotted lines represent the bone boundaries. Scale bar = 50 µm. (12 SL: ColI: *fgfr3^+/+^* n=3; *fgfr3^lof/lof^* n=5; ColVI: *fgfr3^+/+^* n=7; *fgfr3^lof/lof^* n=5; ColXII: *fgfr3^+/+^* n=3; *fgfr3^lof/lof^* n=3) (20 SL: ColI: *fgfr3^+/+^* n=3; *fgfr3^lof/lof^* n=3; ColXI: *fgfr3^+/+^* n=5; *fgfr3^lof/lof^* n=5; ColXII: *fgfr3^+/+^* n=3; *fgfr3^lof/lof^* n=3). (E) Coronal sections of 9, 12 and 20 SL fish imaged by transmission electron microscopy. Scale bar = 0.5 µm. (F) Measurement of fibril transverse area distribution within the suture at 19, 12, and 20 SL revealed a more heterogeneous fibril size in *fgfr3^lof/lof^* compared to controls. (G) Comparison of the mean fibril transverse area in the metopic suture of *fgfr3^+/+ and lof/lof^* fish revealed an increase in fibril size in the mutants at 20 SL (9 SL: (*fgfr3^+/+^* n=3; *fgfr3^lof/lof^* n=2) (12 SL: (*fgfr3^+/+^* n=3; *fgfr3^lof/lof^* n=3) (20 SL: (*fgfr3^+/+^* n=2; *fgfr3^lof/lof^* n=2). F: Frontal bone. Data are presented as mean ± SD. The p-value was determined by a Student’s t test: p < 0.05 = *.

Noting defects in collagen fibrils at the suture, and bone thickening near sutures, we examined whether collagen fibers in the frontal bones were also affected by the absence of Fgfr3 over time. We performed polarization-resolved SHG microscopy at 12 and 26 SL to study collagen fibers structure and organization, assessing the signal intensity and the circular standard deviation of the collagen orientation distribution in a region close to the suture (zone 1) and another farther away (zone 2) (Figure 3A). The circular standard deviation of the collagen orientation distribution measures the degree of variability in the orientation of collagen fibers, providing insight into their alignment and organization at the scale of the image. By 12 SL, no defects in either the structure or the organization were observed in collagen fibers in both regions (Figure 3B and C). In contrast, by 26 SL, although no defect in the collagen structure was noted, a significant disorganization of collagen fibers was observed only in the region adjacent to the suture (zone1) (26 SL *fgfr3^+/+^*: 24.96 ± 4.39 degrees; *fgfr3^lof/lof^*: 31.62 ± 4.13 degrees; p=0.028) (Figure 3B and D). The absence of structural defects and the progressive, localized disorganization of bone collagen fibers at the bone ends suggest that this defect is not directly due to the absence of Fgfr3 but may instead be secondary to changes in suture shape.

**Figure 3.**
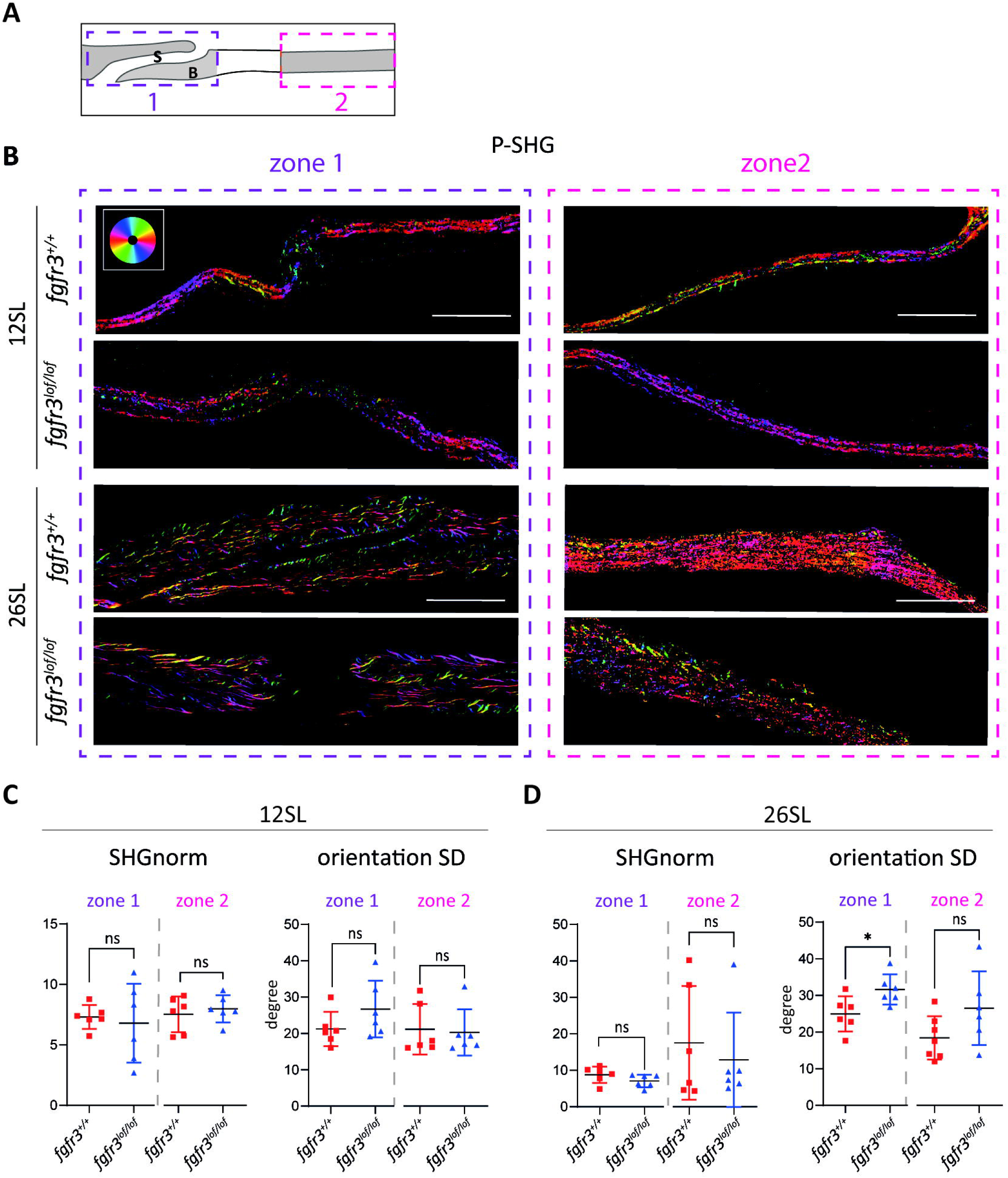
Loss of Fgfr3 affected the organization of bone collagen fibers at the bone ends over time. (A) Diagrams representing the different areas analysed by SHG microscopy. Zone 1 corresponds to the bone surrounding the suture called the bones ends, while zone 2 refers to bone located farther away. (B) Coronal sections of metopic suture at 12 and 26 SL imaged by SHG microscopy at 12 and 26 SL. Scale bar = 50 µm. The color codes for the collagen orientation as shown by the color wheel in the inset. (C-D) Quantification of the SHG intensity normalized to the excitation power squared (SHG_norm_) and of the circular standard deviation of the collagen orientation distribution (orientation SD) in zone 1 and 2 at 12 SL (C) and 26 SL (D). In zone 1, the circular standard deviation of the collagen orientation distribution is significantly higher in *fgfr3^lof/lof^* at 26 SL, suggesting a disorganization of the collagen fibers at the ends of the frontal bones (12 SL: *fgfr3^+/+^* n=6; *fgfr3^lof/lof^* n=6) (26 SL: *fgfr3^+/+^* n=6; *fgfr3^lof/lof^* n=6).Data are presented as mean ± SD. The p-values were determined by Student’s t tests: ns: not significant; p < 0,05 = *.

### The absence of Fgfr3 results in the excessive expansion of osteoprogenitors and inhibits the maturation of osteoblasts at the suture edge

To rule out the hypothesis that excessive bone resorption at the suture edge could be responsible for the change in the shape of the suture, we first assessed osteoclast activity using tartrate-resistant acid phosphatase (TRAP) staining at 12, 20 and 26 SL. No TRAP positive cells were observed in the suture of either *fgfr3^+/+^* or *fgfr3^lof/lof^* fish at the stages tested (Figure 4 supplement 1). This confirmed previous data indicating that osteoclasts are not involved in suture homeostasis in zebrafish, and rules out the possibility that the suture abnormalities observed in our model result from increased osteoclast activity ^30^. We then further explored osteogenesis at the metopic suture from 12 SL to adulthood in *Tg(runx2:GFP; fgfr3^lof/lof or +/+^)* and *Tg(sp7:*mCherry*; bglap:*GFP*; fgfr3^lof/lof or +/+^)* lines. Regarding osteoprogenitors, at 12 SL, *runx2*:GFP+ cells in *fgfr3^+/+^* fish were localized only at the tips of the formed bones, known as the OF. In contrast, in *fgfr3^lof/lof^* fish, *runx2:*GFP+ cells were not only distributed at the tips of the bones but also along the bone surface (Figure 4A). At 20 SL, whereas *runx2*:GFP+ cells in *fgfr3^+/+^* fish were still present only at the tip of the bones, *runx2*:GFP+ cells in *fgfr3^lof/lof^* fish were distributed along the entire edge of the sutures and are no longer present on the bone surface (Figure 4A). This was confirmed by the increased percentage of *runx2*:GFP+ cells present at the edge of the *fgfr3^lof/lof^* sutures compared to controls (*fgfr3^+/+^*: 44.9 ± 24.4% ; *fgfr3^lof/lof^*: 72.1 ±12.5% ; p=0.04) (Figure 4C). Given that osteoprogenitors are characterized by a round shape, we analyzed nuclear circularity of cells present along the suture (Figure 4 supplement 2A-D). For the control, we separately analyzed the nuclei of cells at the OF and those at the edge of the sutures. As expected, in *fgfr3^+/+^* fish, OF nuclei were significantly more circular than those along the suture (Figure 4 supplement 2D). In *fgfr3^lof/lof^* fish, the nuclei along the suture resembled the circularity of OF nuclei in *fgfr3^+/+^* fish (Figure 4 supplement 2D). The modification of shape nuclei was not observed along the bone surface (Figure 4 supplement 2B). These findings were further confirmed by TEM (Figure 4 supplement 2C) and suggests the presence of extra progenitors (*runx2* positive cells) along the suture of *fgfr3^lof/lof^* fish.

**Figure 4.**
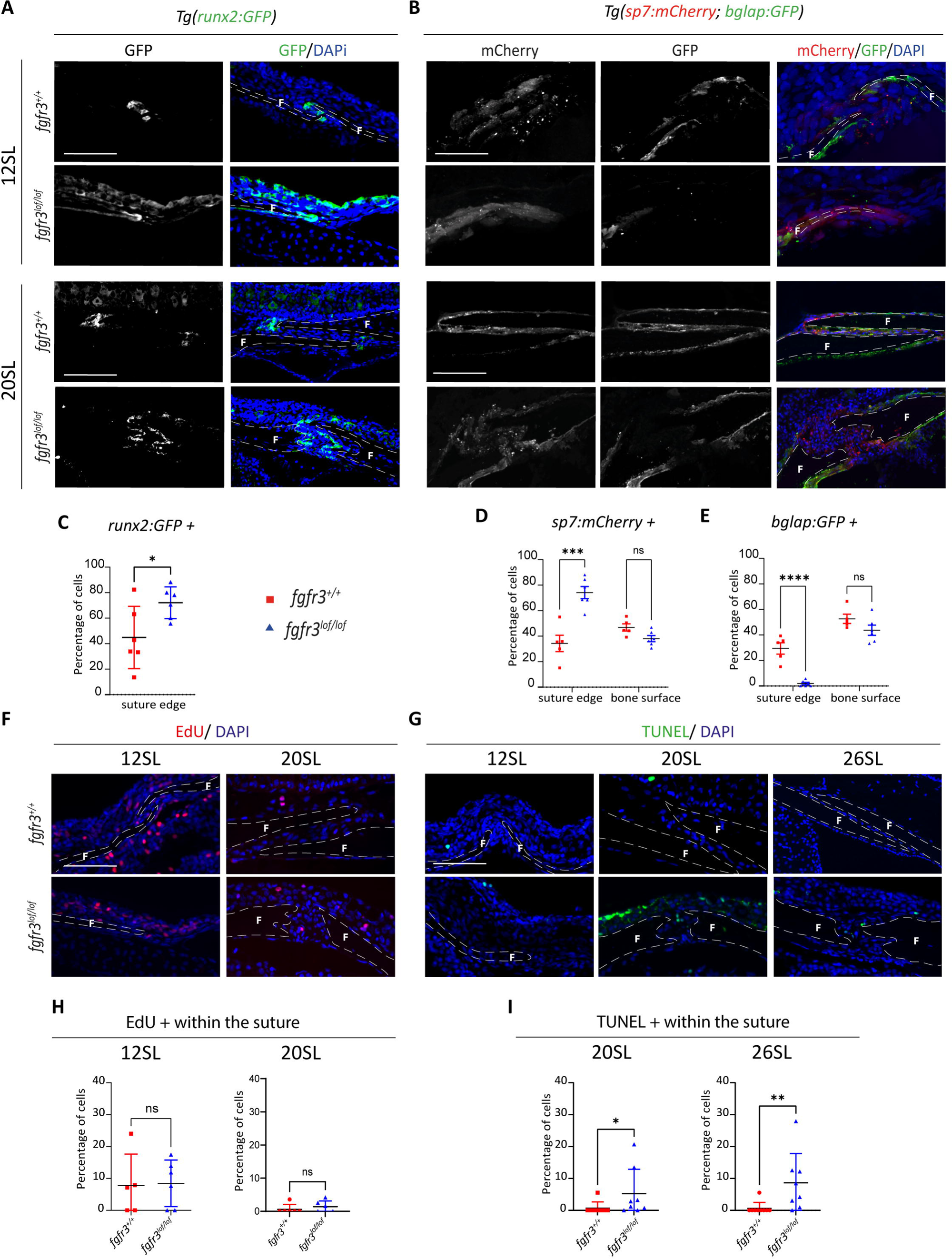
Fgfr3 prevents osteoprogenitor expansion, and activates osteoblast maturation within the metopic suture. (A) Immunofluorescence against GFP (green) performed on metopic sutures of *Tg(runx2:GFP*; *fgfr3^+/+ or lof/lof^*) fish at 12 and 20 SL. Scale bar = 50 µm (B) Immunofluorescence against GFP (green) and mCherry (red) was performed on metopic sutures of *Tg(sp7:mCherry; bglap:GFP*; *fgfr3^+/+ or lof/lof^*) fish at 12 and 20SL. Scale bar = 50 µm. (C) Quantification at 20 SL of the percentage of osteoprogenitors along the suture revealed a significantly higher number of osteoprogenitors in the *fgfr3^lof/lof^* suture compared to the control (*fgfr3^+/+^* n=6; *fgfr3^lof/lof^* n=6). The p-value was determined by a two-way ANOVA. (D-E) Quantification at 20 SL of the percentage of *sp7*:mCherry positive cells and *bglap*:GFP positive cells along the suture and the bone surface revealed a greater number of immature osteoblasts and a drastically lower number of mature osteoblasts at the edge of the *fgfr3^lof/lof^* metopic suture compared to controls. No significant difference was observed along the bone surface. (*fgfr3^+/+^* n=5; *fgfr3^lof/lof^* n=6). The p-values were determined by a two-way ANOVA. (F) EdU assays performed on *fgfr3^+/+ or lof/lof^* metopic suture at 12 and 20 SL. (G) TUNEL assays labelling apoptotic cells performed on *fgfr3^+/+^* and *fgfr3^lof/lof^* metopic suture at 12, 20 and 26 SL fish. Scale bar = 50 µm. (H) Quantification of the percentage of EdU positive cells revealed no significant defect of proliferation at 12 and 20 SL on *fgfr3^lof/lof^* sutures (12 SL: *fgfr3^+/+^* n=5; *fgfr3^lof/lof^* n=6; 20 SL: *fgfr3^+/+^* n=6; *fgfr3^lof/lof^* n=6). p-values were determined by Student’s t tests. (I) Quantitative analysis of apoptotic cell percentages, relative to the number of nuclei stained with DAPI, revealed a significantly higher incidence of apoptosis in the *fgfr3^lof/lof^* suture compared to *fgfr3^+/+^* at 20 and 26 SL (20 SL: *fgfr3^+/+^* n=8; *fgfr3^lof/lof^* n=8; 26 SL: *fgfr3^+/+^* n=9; *fgfr3^lof/lof^* n=8). p-values were determined by Student’s t tests. Nuclei were counterstained with DAPI (blue) and the dotted lines represent the bones boundary. F: Frontal bone. Data are presented as mean ± SD. ns: not significant; p < 0,05 = *; p < 0,01 = **; p < 0,001 = ***; p < 0,0001 = ****.

Next, we studied the impact of the absence of Fgfr3 on immature (*sp7-positive*) and mature (*bglap-positive*) osteoblasts. In both *fgfr3^+/+^* and *fgfr3^lof/lof^* fish, immature osteoblasts were present throughout the bone surface and along the suture at all stages (Figure 4B and Figure 4 supplement 3A). The quantification of immature osteoblasts at 20 and 26 SL showed significantly more immature osteoblasts at the edge of the metopic suture in the *fgfr3^lof/lof^* model compared to controls, with no difference along the bone surface (suture, *fgfr3^+/+^*: 34.3 ± 14.3%; *fgfr3^lof/lof^*: 74.1 ± 11.7%; p<0.0001; bone surface, *fgfr3^+/+^*: 46.7 ± 6.3%; *fgfr3^lof/lof^*: 38.0 ±5.9%; p=0.315) (Figure 4D and Figure 4 supplement 3B). Regarding mature osteoblasts, in *fgfr3^+/+^* fish we observed their presence throughout the bone surface and along the suture at all stages (Figure 4E and Figure 4 supplement 3C). Interestingly, in *fgfr3^lof/lof^,* while mature osteoblasts were present throughout the bone surface similar to controls, there was a near absence of mature osteoblasts within the suture at all stages (suture, *fgfr3^+/+^*: 29.4 ± 10%; *fgfr3^lof/lof^*: 1.9 ± 2.4%; p<0.0001; bone surface, *fgfr3^+/+^*: 52.7 ± 8.1%; *fgfr3^lof/lof^*: 43.7 ± 9.8%; p=0.1586). To determine whether these events occur in other sutures, we analyzed immature and mature osteoblasts in the coronal suture at 26 SL(Figure 4 Supplement 3D-F). In *fgfr3^lof/lof^* coronal sutures, a higher number of immature osteoblasts (*fgfr3^+/+^*: 46.2 ± 6.6%; *fgfr3^lof/lof^*: 75.3 ± 5.9%; p=0.0011) and an absence of mature osteoblasts (*fgfr3^+/+^*: 50.9 ± 20.6%; *fgfr3^lof/lof^*: 0 ± 0%; p<0.0001) were found compared to controls, with no significant difference for the immature and mature osteoblasts along the bone surface (*sp7*: 26 SL, *fgfr3^+/+^*: 60.8% ± 17.8%; *fgfr3^lof/lof^*: 73.9% ± 7.5%; p=0.1355; *bglap*: 26 SL, *fgfr3^+/+^*: 47.8% ± 17.6%; *fgfr3^lof/lof^*: 34.6% ± 7.6%; p=0.3450). We conclude that the same osteoblast maturation abnormalities are present in *fgfr3^lof/lof^* coronal sutures as in the metopic suture.

To better understand the higher number of *runx2* and *sp7* positive cells as well the absence of *bglap* positive cells within the suture of the *fgfr3^lof/lof^* fish, we first studied cell proliferation rates at 12 and 20 SL by performing balneation EdU experiments. Surprisingly, we found no difference at either stage (Figure 4F, H) suggesting that the absence of Fgfr3 does not lead to a proliferation defect of osteoprogenitors and immature osteoblasts within the suture. Subsequently, we investigated apoptosis by performing a TUNEL assay. At 12 SL, we observed no apoptosis for either genotype. However, at 20 and 26 SL, a few randomly localized apoptotic events were observed only within the *fgfr3^lof/lof^* suture (20 SL; *fgfr3^+/+^*: 0.7%; *fgfr3^lof/lof^*: 5.2%; p=0.0294; 26 SL; *fgfr3^+/+^*: 0.6%; *fgfr3^lof/lof^*: 8.7%; p=0.0019) (Figure 4G, I). Since apoptosis appears to occur randomly and later than the observed absence of mature osteoblasts, it does not account for their disappearance. The latter is likely due to the inhibition of maturation of immature osteoblasts.

All these data demonstrate that the absence of Fgfr3 seems to have a dual effect specifically within the cranial sutures. On one hand, it induces the expansion of osteoprogenitors (which normally localize only to the tip of the bones) all along the suture edge, which then differentiate into immature osteoblasts. On the other hand, it inhibits immature osteoblasts maturation, leading to their accumulation at the suture edge.

### The absence of Fgfr3 alters the SuSC marker profile, highlighting the downregulation of the BMP antagonist Grem1 and the activation of the Wnt signaling pathway within the suture

Given the greater number of osteoprogenitors in the *fgfr3^lof/lof^* sutures compared to the control (Figure 4D), we also studied the expression of *prrx1, gli1, axin2,* and *grem1* at 12 and 20 SL. All these genes are known to be expressed in distinct populations of SuSCs, and play a crucial role in maintaining suture patency (Figure 5A-B) ^11,31,5^. *prrx1* was expressed within the suture and bone surface at both stages, with no difference in expression between the genotypes (Figure 5A-B). The expression of *gli1* was nearly absent at 12 and 20 SL in *fgfr3^+/+^* sutures, and at 20 SL in *fgfr3^lof/lof^* sutures. We only observed a slightly higher expression of *gli1* at 12 SL in *fgfr3^lof/lof^* sutures compared to controls (Figure 5A-B), but this is unlikely to explain the increased number of cells within the suture and may instead reflect only the delay of the mutant cranial vault growth.

**Figure 5.**
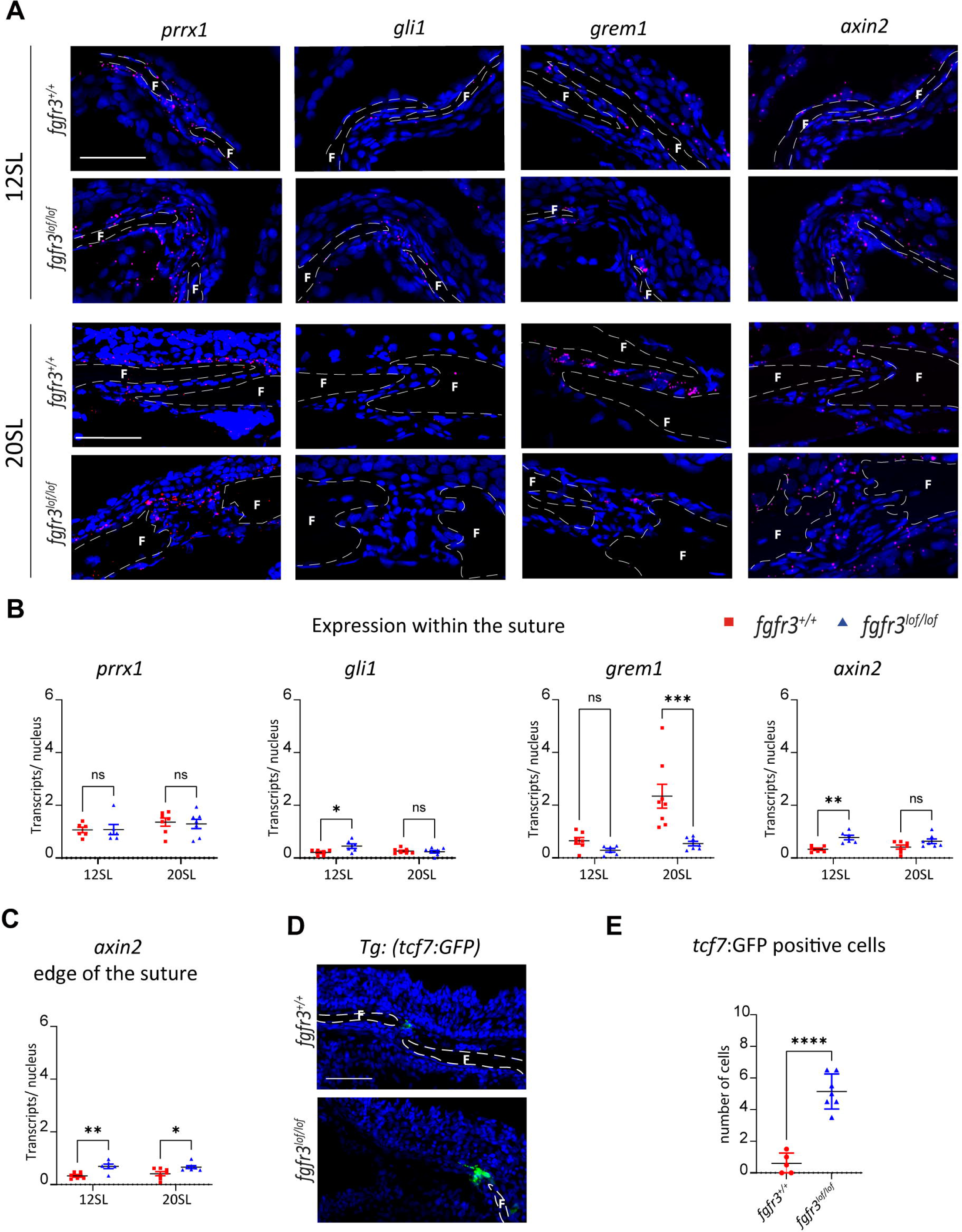
The absence of Fgfr3 alters the SuSC marker profile, highlighting the downregulation of the BMP antagonist Grem1 and the activation of the Wnt signaling pathway within the suture. (A) Expression of *prrx1, gli1, axin2*, and *grem1* (red) in coronal sections of the metopic suture at 12 and 20 SL, labeled by RNAscope in situ hybridization. Nuclei are counterstained with DAPI (blue). Scale bar = 50 µm. (B) Quantitative RNAscope analysis revealed increased expression of *gli1* and *axin2* in *fgfr3^lof/lof^* metopic sutures compared to controls. In contrast, *grem1* expression is significantly reduced at 20 SL. (12 SL: *fgfr3^+/+^* n=6; *fgfr3^lof/lof^* n=6; 20 SL: *fgfr3^+/+^* n=7; *fgfr3^lof/lof^* n=7). p-values were determined by two-way ANOVA. (C) Quantitative RNAscope analysis of *axin2* expression along the metopic suture edge demonstrated higher *axin2* expression in *fgfr3^lof/lof^* fish compared to controls at both 12 and 20 SL (12 SL: *fgfr3^+/+^* n=6; *fgfr3^lof/lof^* n=6; 20 SL: *fgfr3^+/+^* n=7; *fgfr3^lof/lof^* n=7). p-values were determined by two-way ANOVA. (D) Immunofluorescence against GFP (green) was performed on metopic sutures of Tg(7xTCF-Xla.Siam:GFP; *fgfr3^+/+ or lof/lof^*) fish at 12 SL. (E) Quantification at 12 SL of the number GFP-positive cells revealed a significantly higher number of cells at the tips of *fgfr3^lof/lof^* bones compared to controls, suggesting an overactivation of the Wnt signaling pathway (*fgfr3^+/+^* n=5; *fgfr3^lof/lof^* n=7). p-values were determined by Student’s t-test: ns = not significant; p < 0.05 = *; p < 0.01 = **; p < 0.001 = ***; p < 0.0001 = ****. Data are presented as mean ± SD.

The expression of *grem1,* which encodes Gremlin1, an antagonist of BMP, was restricted to the suture and increased from 12 to 20 SL in controls. In contrast, in *fgfr3^lof/lof^,* its expression at 12 SL was restricted to the tips of the bones, and was significantly lower at 20 SL compared to controls (20 SL: *fgfr3^+/+^*: 2.3 ±1.2 transcripts/nucleus (t/n); *fgfr3^lof/lof^*: 0.5± 0.3 t/n; p=0.0001) (Figure 5A-B).

However, we observed a significant upregulation of *axin2* in mutants compared to controls at 12 SL within the suture, which appeared to be maintained over time, although not significantly (12 SL: *fgfr3^+/+^*: 0.3 ±0.1 t/n; *fgfr3^lof/lof^*: 0.8± 0.2 t/n; p=0.0017; 20 SL: *fgfr3^+/+^*: 0.4 ±0.3 t/n; *fgfr3^lof/lof^*: 0.6± 0.2 t/n; p=0.09). Given that *axin2* seemed to be more expressed at the edges of the suture, where osteoprogenitors and immature osteoblasts are localized, we analysed its expression specifically at the suture edge (Figure 5C). We found higher expression of *axin2* at both 12 and 20 SL in *fgfr3^lof/lof^* suture edges compared to controls (12 SL: *fgfr3^+/+^*: 0.3 ±0.1 t/n; *fgfr3^lof/lof^*: 0.7± 0.2 t/n; p=0.004; 20 SL: *fgfr3^+/+^*: 0.4 ±0.2 t/n; *fgfr3^lof/lof^*: 0.7± 0.1 t/n; p=0.03). As Axin2 is both a Wnt/β-catenin target gene and a negative regulator of the Wnt canonical pathway, we aimed to determine whether there is abnormal activation of the Wnt signaling pathway at the edge of *fgfr3^lof/lof^* metopic sutures ^32^. To do this, we used the Wnt reporter line *Tg(7xTCF-Xla.Siam:GFP)ia4* to generate *Tg(7xTCF-Xla.Siam:GFP)^ia4^; fgfr3^lof/lof or +/+^* fish and examined the presence of GFP-positive cells at the metopic suture at 12 SL. We observed that GFP positive cells were localized at the tips of the frontal bones in both genotypes but a greater number of positive cells were present in *fgfr3^lof/lof^* fish compared to controls, indicating activation of the Wnt pathway (*fgfr3^+/+^*: 0.6 ±0.6; *fgfr3^lof/lof^*: 5.1± 1.1; p<0.0001) (Figure 5D and E).

The interplay between FGF and Wnt signaling in cranial sutures was recently highlighted by Bobzin et al., who showed in a FGFR2 LoF mouse model that FGF signaling regulates anterior fontanelle closure in mice by inhibiting Wnt signaling, guiding Scx- and Tnmd-expressing ectocranial mesenchyme toward bone and cartilage differentiation ^33^. Interestingly, *tnmd* expression was found downregulated in *fgfr3* LoF zebrafish during cranial bone development ^28^. Therefore, we investigated whether Fgfr3 LoF and Wnt pathway activation influenced the expression of *tnmd* or *scx* within the suture at 20 SL. Contrary to findings in mice, neither gene was dysregulated nor restricted to a specific cell population in either the control or *fgfr3^lof/lof^* suture (Figure 5 supplement 1). These findings suggest that the mechanisms leading to suture anomalies differ between the animal models. Nevertheless, our data indicate that the absence of Fgfr3 induces overactivation of Wnt signaling at the suture edge and potentially of the BMP pathway within the suture.

### The absence of Fgfr3 is not compensated by other Fgfrs and leads to the overexpression of Fgf18 within the suture

Our data reveal that Fgfr3 has a localized effect within the suture, which becomes particularly evident in adulthood (from 20 SL) with the presence of ectopic osteoprogenitors and the absence of mature osteoblasts exclusively at the suture edges, while osteogenesis along the bone surface appears unaffected. These results were unexpected, as *fgfr3* expression was observed both within the sutures and along the bone surface (Figure 1B and C). This differential activity of Fgfr3 may reflect the varying expression levels of its ligands, such as Fgf2 and Fgf18, which are known to be expressed during zebrafish cranial vault development ^28^. Therefore, we examined the expression of *fgf2* and *fgf18* (Figure 6A). With regard to *fgf2*, no differential expression was observed between the bone surface and the suture, regardless of the genotype at 12 and 20 SL (Figure 6B and C). However, in the case of *fgf18*, although its expression was low in both the bone surface and suture in *fgfr3^+/+^* fish, we observed higher expression of *fgf18* in *fgfr3^lof/lof^* sutures compared to the *fgfr3^+/+^* sutures at both stages (12SL: *fgfr3^+/+^*: 0.3 ±0.1 t/n; *fgfr3^lof/lof^*: 0.7± 0.2 t/n; p=0.004; 20 SL: *fgfr3^+/+^*: 0.4 ±0.2 t/n; *fgfr3^lof/lof^*: 0.7± 0.1 t/n; p=0.03) (Figure 6B and C). The absence of differential expression of *fgf2* and *fgf18* between the bone surface and suture in *fgfr3^+/+^* fish may suggest that the localized role of Fgfr3 within the suture is not dependent on the expression of these two ligands. However, these results highlight that the absence of Fgfr3 leads to an increase of *fgf18* expression specifically within the suture.

**Figure 6.**
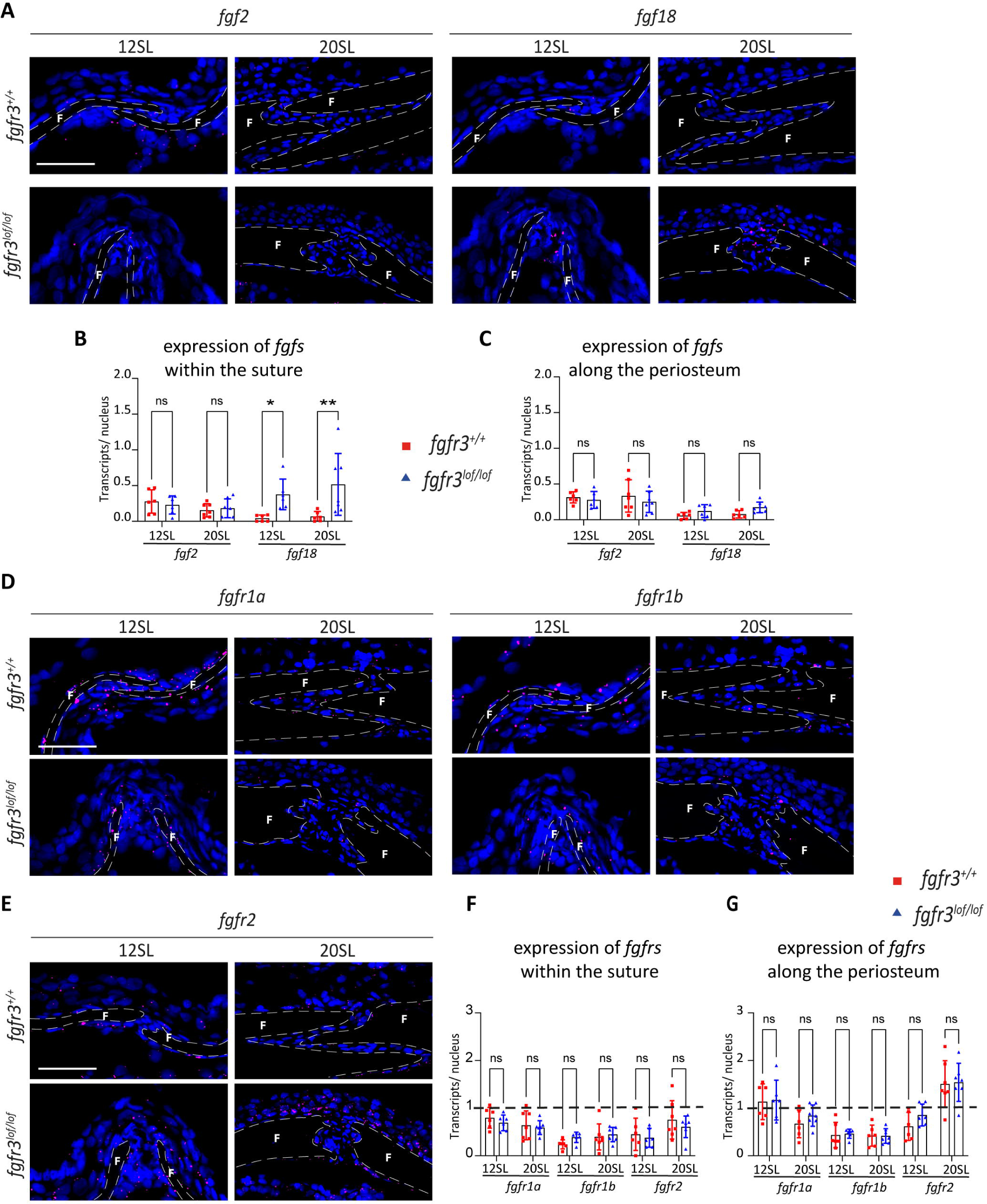
The absence of Fgfr3 is not compensated by other Fgfrs and leads to the overexpression of Fgf18 within the suture. (A) *fgf2* and *fgf18* expression (red) on coronal sections of metopic suture at 12 and 20 SL labelled by RNAscope *in situ* hybridization. Nuclei were counterstained with DAPI (blue). Scale bar = 50 µm. The dotted lines represent the bone boundaries. (B-C) RNAscope Quantitative analyses of *fgf2* and *fgf18* expression in *fgfr3^+/+^* versus *fgfr3^lof/lof^* within the suture (B) or along the bone surface (C) at 12 and 20 SL showed that *fgf18* is significantly more expressed in the *fgfr3^lof/lof^* suture than in *fgfr3^+/+^* suture at 12 and 20 SL (12 SL and 20 SL: *fgfr3^+/+^* n=6; *fgfr3^lof/lof^* n=6). (D, E) *fgfr1a, fgfr1b* and *fgfr2* expression (red) on coronal sections of metopic suture at 12 and 20 SL labelled by RNAscope *in situ* hybridization. Nuclei were counterstained with DAPI (blue). Scale bar = 50 µm. The dotted lines represent the bone boundaries. (F, G) RNAscope quantitative analyses of *fgfr1a, fgfr1b* and *fgfr2* expression in *fgfr3^+/+^* versus *fgfr3^lof/lof^* within the suture (F) or along the bone surface (G) at 12 and 20 SL revealed similar expression of *fgfr1a, fgfr1b*, and *fgfr2* in both genotypes. (12 SL: *fgfr3^+/+^* n=6; *fgfr3^lof/lof^* n=6; 20 SL: *fgfr3^+/+^* n=7; *fgfr3^lof/lof^* n=7). Data are presented as mean ± SD. p-values were determined by two-way ANOVA: ns: not significant; p < 0,05 = *; p < 0,01 = **.

We next examined the expression of *fgfr1a*, *fgfr1b* and *fgfr2* (Figure 6D-G). *fgfr1b* was consistently expressed at a low level throughout cranial suture formation, with no significant variation observed based on its spatial distribution or genotype. The expression of *fgfr1a* and *fgfr2* also did not differ between control and mutant. But when we compared Fgfr2 expression in suture vs bone surface, we observed that *fgfr2* is significantly more expressed in both the *fgfr3^+/+^* and *fgfr3^lof/lof^* bone surface at 20 SL (*fgfr3^+/+.^* bone surface: 1.5 ±0.5 t/n; suture: 0.8± 0.4 t/n; p=0.004; *fgfr3^lof/lof^*; bone surface: 1.5 ±0.4 t/n; suture: 0.6± 0.2 t/n; p<0,0001) suggesting a more prominent role for *fgfr2* in the bone surface (Figure 6 supplement 1). These results revealed, first, that neither the absence of Fgfr3 nor the increased expression of Fgf18 leads to an increase in expression of other Fgfrs; second, that other Fgfrs may play a more prominent role than Fgfr3 at the bone surface; and third, the key role of Fgfr3 for proper suture formation in zebrafish.

## Discussion

By examining cranial sutures from initiation to adulthood in our LoF Fgfr3 zebrafish model, we demonstrated for the first time that Fgfr3 is essential for the proper formation of cranial sutures (Figure 7) ^28^. We showed that the absence of bone overlap in *fgfr3^lof/lof^* cranial sutures was linked to an increased number of cells surrounded by an ECM, with defects in collagen fibrilllogenesis. In addition, we observed a progressive bone thickening accompanied by disorganization of collagen fibers at their tips. Furthermore, we provide evidence that Fgfr3 is crucial for limiting the number of osteoprogenitors at the bone ends and for promoting the maturation of osteoblasts specifically at the suture edge. Finally, our results suggest that Fgfr3 regulation of cranial suture formation involves both the canonical Wnt and BMP signaling pathways and highlight a potential regulatory loop through the modulation of *fgf18* expression. These results are essential as until now, the involvement of FGFR3 in cranial suture formation had only been illustrated by the existence of the craniosynostoses associated with FGFR3 mutations (Muenke and CAN syndrome) and a few studies reporting occasional premature fusion of coronal sutures in mouse models expressing Fgfr3 GoF mutations ^19,20,23,34^.

**Figure 7.**
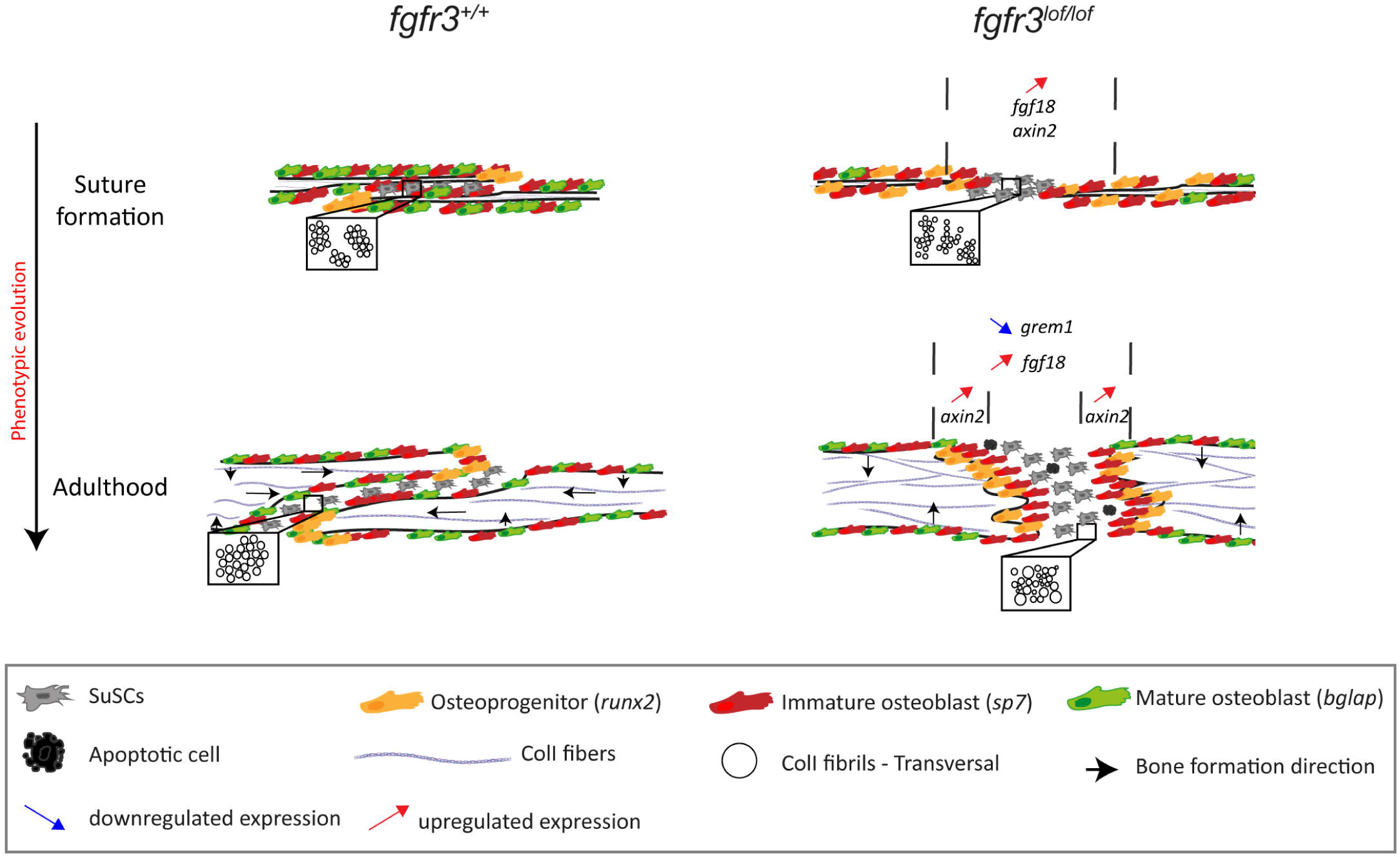
Impact of Fgfr3 absence on cranial suture formation. The absence of FGFR3 exerts pleiotropic effects on cranial suture formation, leading to significant structural defects. Without Fgfr3, proper overlap of the sutures is prevented, resulting in bone widening. These structural abnormalities are associated with osteogenesis defects, as Fgfr3 deficiency disrupts the expansion of osteoprogenitors within the suture and hinders bone maturation, thereby inhibiting longitudinal bone growth. Additionally, the absence of Fgfr3 affects the extracellular matrix (ECM) at the ends of bones and within the suture itself. From a signaling pathway perspective, the loss of Fgfr3 triggers the overexpression of Fgf18 and causes the misregulation of *axin2* and *grem1* expression. This positions Fgfr3 as a central regulator of cranial suture formation in a complex network involving FGF, BMP, and Wnt signaling pathways.

Our study focuses on the late stages of cranial vault development, when the bones converge and the suture forms. In this phase, periosteal osteoblasts drive skull widening and remodeling, while suture osteoblasts are responsible for maintaining continuous cranial expansion and keeping the suture open. Analysis of osteogenesis at both sites using *Tg(runx2:GFP; fgfr3^lof/lof or +/+^),* and *Tg(sp7:*mCherry*; bglap:*GFP*; fgfr3^lof/lof or +/+^)* lines showed that Fgfr3 loss does not affect osteogenesis along the bone surface. In contrast, from the suture initiation onwards, at bone extremities, Fgfr3 absence increases the number of osteoprogenitors (Runx2-positive cells), which are no longer confined to bone tips, leading to an expanded ossification front over time. Interestingly, this increase of Runx2 positive cells associated with the lack of bone overlap has been described in *sp7*^-/-^ fish ^35^. In that study, the authors linked the increase in Runx2 positive cells to the absence of *sp7* expression, which is essential for the normal progression of osteoblast differentiation. In contrast, in *fgfr3^lof/lof^* fish, differentiation appears blocked not at the transition from osteoprogenitors to immature osteoblasts but from immature to mature osteoblasts. This is evidenced by the higher number of *sp7^+^* cells and the absence of *bglap^+^* cells at the suture edge, unrelated to apoptosis. These findings showed that Fgfr3 positively regulates osteoblast maturation during suture formation, as we demonstrated at an earlier stage of cranial bone development ^28^. However, the increase in *runx2-* and *sp7*-positive cells remains complex to interpret, as no increase in osteoprogenitor or immature osteoblast proliferation was detected at the suture edge. Instead, it may result from enhanced recruitment of osteoprogenitors from differentiating SuSCs and/or the accumulation of cells that fail to mature. In any case, this osteogenesis defect, localized at bone extremities but not along the bone surface, likely underlies both the progressive widening of the bone extremities and a part of the increased cellularity observed within the suture.

Our work reveals that the absence of Fgfr3 leads to changes in the expression of *grem1* and *axin2*, which are both known to be expressed in SuSCs and are involved respectively, in the BMP and canonical Wnt signaling pathways, two major pathways crucial for cranial suture homeostasis and osteoblast differentiation ^36^. First, we observed a significant decrease in *grem1* expression within the sutures in the absence of Fgfr3. Grem1 is an antagonist of the BMP signaling pathway recently identified as a marker of a SuSC subpopulation in zebrafish ^5^. Interestingly, Farmer et al., reported that the absence of *grem1* in zebrafish leads to a slight delay of cranial vault formation. However, when Grem1 is absent along with Nog2 and Nog3, two other BMP antagonists, it leads to overactivation of BMP signaling in the suture, resulting in the same suture anomalies seen in our *fgfr3^lof/lof^* model (e.g., failure of bone overlaps and the presence of ectopic bone). This phenotypic similarity and the decreased *grem1* expression in *fgfr3^lof/lof^* metopic sutures suggest that the absence of Fgfr3 may increase BMP signaling within the suture. Notably, BMP pathway activation was also reported in the *sp7^-/-^* zebrafish model further supporting our hypothesis ^35^. As the BMP pathway promotes osteogenesis, its activation in the absence of Fgfr3 may increase osteoprogenitor recruitment along the suture. Further experiments are required to validate this hypothesis and to determinate the link between the absence of Fgfr3 and the decrease of *grem1* expression. Furthermore, existing data on cranial sutures suggest the opposite that FGF signaling promotes BMP signaling ^37,38^. Additionally, it was reported in mouse model that FGF18 modulates *Grem1* expression via the activation of FGFR3 in chondrocytes located in synchondroses or in articular cartilage ^39,40^. Interestingly we noted in absence of Fgfr3 an overexpression of *fgf18*, one of the Fgfr3 ligands essential for cranial vault development^41^. Although regulation of Grem1 by Fgf18 in cranial sutures has not been documented, we could hypothesize that Fgf18 regulates *grem1* expression through its binding to other Fgfrs. At the RNA level, we did not observe an increase in *fgfr1a*, *fgfr1b*, and *fgfr2* expression within the suture in the absence of Fgfr3, suggesting the absence of a compensatory mechanism. Nevertheless, since *fgfr1a, fgfr1b, and fgfr2* are expressed within the suture, albeit to a lesser degree than Fgfr3, their activation through Fgf18 binding remains a possibility.

Additionally, we showed that the absence of Fgfr3 appears to lead to overactivation of the Wnt signaling pathway at the suture edges, as evidenced by both the overexpression of *axin2*, a Wnt/β-catenin target gene and the increased number of *7xTCF-Xla.Siam:*GFP-positive cells in Fgfr3 deficient fish. This suggests that Fgfr3 may act as a potential inhibitor of the canonical Wnt signaling pathway. Notably, Fgfr3 has already been described as such during chondrogenesis, specifically in pharyngeal skeleton formation in zebrafish ^27^. Moreover, recent work by Bobzin et al. highlighted crosstalk between FGF and Wnt/β-catenin pathways in sutures of mice, demonstrating that FGFR2 activation at the OF of the frontal bones regulates anterior fontanel closure by inhibiting Wnt signaling through Wif1 ^33^. Within the suture, Wnt signaling is involved in regulating the number of SuSCs and also osteoblast differentiation ^42^. Aberrant β-catenin activation has been shown to widen all cranial sutures in mice, accompanied by increased *Runx2* and *Sp7* expression, the absence of the mature osteoblast marker *Osteopontin*, and enhanced BMP signaling ^43^. These findings align with our observations and further support that Fgfr3 might promote maturation of osteoblasts via the inhibition of canonical Wnt signaling in cranial sutures.

Notably*, Fgf18* is a direct Wnt target gene, regulated positively via a TCF/Lef binding site, in cooperation with RUNX2 ^44^. Thus, Wnt pathway activation may drive *fgf18* overexpression in the absence of Fgfr3, thereby outlining the framework of the interplay between the Fgfr3, Wnt, and BMP pathways.

Finally, our work provides evidence that the absence of Fgfr3 may impact osteogenesis not only through direct pathway activation but also by altering the microenvironment of the cells, particularly the collagen network, which, by transmitting biomechanical signals influences cell fate. In our previous study, we showed that Fgfr3 loss leads to the overexpression of ECM-regulating proteins like *col6a1, col12a1a,* and *col12a1b* (Dambroise et al., 2020). Collagen XII is a FACIT collagen known to bind to collagen I fibrils, contributing to the collagen suprastructure and helping to absorb biomechanical stress ^45^. We show here that the loss of Fgfr3 disrupts the organization of ColI fibrils in the bones near the suture and within the suture itself, due to both a delay in cranial suture formation and a modification of its shape. This is accompanied by a fibrillogenesis defect, potentially linked to ColXII overexpression, as ColXII regulates both collagen maturation and organization ^15,46^. We suggest that modifications in the osteogenic cell environment, including collagen network remodeling and the predominance of lateral over longitudinal bone growth, may alter the mechanical forces exerted on osteogenic cells within the suture. These changes could influence cell fate and potentially explain the progressive emergence of random apoptosis, an as-yet unexplained phenomenon.

In conclusion, our work using a zebrafish model is the first to demonstrate that Fgfr3 is a crucial regulator of cranial suture formation, essential for proper osteogenesis and suture development (Figure 7). We highlight its importance in the regulation of BMP and Wnt pathways within the sutures, the underlying mechanisms of which remain to be validated and further explored. Altogether, these findings open new perspectives for understanding suture homeostasis and FGFR3-related craniosynostoses, paving the way for new targeted therapies.

## Materials and methods

### Zebrafish husbandry, fgfr3^lof^ fish and transgenic lines

Zebrafish were raised, hatched and maintained in an aquatic facility at 28°C (approval number 2018080216094268 and APAFIS #36172-2022041412092374 v1), in accordance with European Directive 2010/63/EU for animals. Developmental stages were determined by the standard length (SL) of the fish, measured from the snout to the caudal peduncle ^47^. *fgfr3^lof^* fish correspond to the *fgfr3^lof1^* line described previously^28^. Transgenic lines *Tg(runX2 :GFP), Tg(sp7:mCherry), Tg(bglap:GFP)* ^48^ and *Tg(7xTCF-Xla.Siam:GFP)^ia4 49^* labelling respectively osteoprogenitors, immature osteoblasts, mature osteoblasts and activation of Wnt signalling have been previously described.

### Genotyping

Genomic DNA was isolated from caudal fin biopsies under anaesthetic with benzocaine (100mg/L). Tissues were lysed in 10 mM Tris-HCl, pH 8.5, 2 mM EDTA, pH 8, 0.2% Triton X-100, and 250 μg/mL proteinase K (Sigma-Aldrich, St. Louis, MO, USA) for at least 1 hr at 55°C. The samples were genotyped in heteroduplex mobility assay using the following primers *fgfr3lof*_F: 5’-TGAACGATCCCAGGAATCCC -3’ and *fgfr3lof*_R: 5’-GGTGAGAATGAAGAGCACGC -3’.

### Histological analyses

Zebrafish at 12, 20 and 26 SL were fixed for 24 hrs in 4% paraformaldehyde at 4°C, the heads were decalcified in a 0.5M EDTA solution pH8 for 1 to 5 days, then embedded in paraffin. Paraformaldehyde-Fixed and Paraffin-Embedded (FFPE) (4µm) sections were cut either transversally or sagittally. Haematoxylin Eosin (HE) and Sirius red-Alcian blue staining were performed after dewaxing with Xylene substitute, Neo-Clear® followed by rehydration with decreasing ethanol baths. For HE staining, the sections were incubated sequentially with Haematoxylin for 30 seconds, LiCO_3_ for 2 seconds and Eosin for 10 seconds. For Sirius Red-Alcian blue staining, sections were incubated in Weigert’s solution (Hematein 0.1g/mL in EtOH 95° and FeCl3 29% HCl 37% v/v) for 5 minutes, washed with acetic acid 1%, incubated 30 min with Alcian blue (1mg/mL; 10% HCl 1N), and finally treated with 0.1% direct red diluted in aqueous saturated picric acid for 2 hours. Image acquisitions were performed using an Olympus IX81 microscope equipped with an Olympus IX2-UCB camera. For HE staining, the sample sizes were n=6 *fgfr3^+/+^* and n=6 *fgfr3^lof/lof^* at 12SL, n=6 *fgfr3^+/+^* and n=8 *fgfr3^lof/lof^* at 20SL and n=10 *fgfr3^+/+^* and n=9 *fgfr3^lof/lof^* at 26SL. Quantification of bone thickness was performed on the two bones of one section per individual using the Fiji software (ImageJ). For Sirius Red-Alcian blue, n=6 *fgfr3^+/+^* and n=6 *fgfr3^lof/lof^* at each stage were analyzed.

### Immunohistochemistry (IHC)

Zebrafish at 12 SL, 20 SL and 26 SL were fixed for 24 hrs in 4% paraformaldehyde at 4°C, heads were decalcified for 1 to 5 days in a 0.5M EDTA solution pH8 and impregnated in sucrose 30% during at least 10 days, then, embedded in frozen medium containing 30% sucrose and 7% gelatine. 20 µm sections were cut in transverse orientations. The polyclonal primary antibodies used are anti-Collagen I (Col1α1, Abcam, ab23730), anti-collagen α1 VI (Col6a1, ^50^), anti-zebrafish collagen XII antibodies (Col12a1, ^51^), anti-mCherry (Takara, 632496) and anti-GFP (Invitrogen; A11122). Initial steps of IHC and blocking buffers depended on the primary antibody used. For ColI labelling, after six 10-minutes washes with PBSTr (Phosphate-Buffered Saline (PBS (Gibco)) Triton-X100 0,5%), the sections were blocked in blocking buffer BB1 (BSA 4%; Triton-X100 0.1%, PBS) for 1hr at room temperature (RT) and then incubated overnight at 4°C with anti-ColI diluted 1:200 in BB1. Before secondary antibody incubations, sections were washed three times for 5 min. For ColXII, mCherry and GFP labelling, the sections were incubated at RT for 1 hr in BB2 (3% sheep serum; 1% BSA in PBS), followed by a 1 hr incubation with the corresponding primary antibody diluted 1:250 in BB2 and then washed three times for 5 min with PBS. For Col6 labelling, after 20 min wash in PBS, sections were blocked in BB3 (Goat serum 5% diluted in PBS) for 30 min, incubated overnight at 4°C with the corresponding primary antibody diluted 1:400 in BB3 and washed three times for 5 min with PBS. The final steps of IHC for whole labelling consist of a 1 hr incubation with an Alexa fluor^TM^ goat anti-rabbit 594 or 647 (Invitrogen) diluted 1:500 in PBS followed by three 5-minutes washes with PBS and mounting of the slides in DAPI Fluoromount-G ^TM^ (Electron Microscopy Sciences -#17984-24). IHC were imaged with a Spinning Disk Zeiss controlled by Zen Blue software (Olympus, Tokyo, Japan) and processed on the Fiji software (ImageJ) to combine all the images into one containing the maximum intensity information for each pixel. The analysis of ColI, ColVI and ColXII, which are not quantitative was assessed at least on n=3 per genotype for each different stage. The mCherry and GFP labelling for the *Tg(runx2 :GFP), Tg(sp7:mCherry), Tg(bglap:GFP)* were quantified for each stage with at least n=5 samples on one section from each individual at 20 and 26 SL per genotype. The quantification represents the ratio between the number of positively labeled cells and the number of DAPI-stained cells.

### EdU and TUNEL labelling

For EdU analyses, 12 and 20 SL*Tg(runX2 :GFP)* fish were immersed in an EdU solution at 12.5 µg/ml during 48hpf. EdU incorporation was subsequently detected on frozen sections (30% sucrose, 7% gelatin) for 12 SL fish and on FFPE sections for 20 SL fish. The EdU was detected using the EdU Click-iT Plus EdU Alexa Fluor 647 Imagin g kit (Life Technologies, Carlsbad, CA, USA), following the manufacturer’s protocol. Sample sizes were n=6 *fgfr3^+/+^* and n=6 *fgfr3^lof/lof^* at each stage. TUNEL assay were performed on the FFPE sections using the DeadEnd Fluorometric TUNEL system (Promega, Madison, WI, USA), following the manufacturer’s instructions (permeabilization step with proteinase K at 10mg/mL diluted in a PBS at 1:500 for 7 min). Images were captured with a Spinning Disk Zeiss controlled by Zen Blue software (Olympus, Tokyo, Japan) and processed using the Fiji software (ImageJ) to combine all images into one containing the maximum intensity information for each pixel. The TUNEL experiment was quantified at 20SL and 26SL for n=8 per genotype on two adjacent sections per individual. At 12SL, TUNEL experiment was performed only on n=3 per genotype due to the absence of positive cells.

### RNAscope Assay

RNAscope Assays were performed on FFPE sections according to the manufacturer’s instructions « RNAscope Multiplex Fluorescent Reagent Kit v2 Assay » provided by ACD a bio-techne brand. After dewaxing, the slides were incubated with Target Retrieval and Protease Plus reagents for 15 minutes and 30 minutes, respectively. The zebrafish *fgfr1a* (C1-409441); *fgfr1b* (C2-409451); *fgfr2* (C1-420961); *fgfr3* (C1-438971); *fgf2* (C1-420951); *fgf8a* (C2-559351); *fgf18* (C3-123572); *axin2* (C1-465351); *gli1* (C2-542721); *prrx1* (C3-535321); *tnmd* (C1-564481); *scx* (C2-564451); *grem1a* (C1-535291) probes were designed and synthesized by the manufacturer. The universal negative control with probes targeting dapB gene (320871) provided by ACD was used as reference. Double labeling was performed following the manufacturer’s instructions. The probes were incubated for 2 hrs at 40°C, with a dilution of 1/50 for each probe. Opal 570 (Akoya, Cat. No. FP1488001KT) and Opal 650 (Akoya, Cat. No. FP1496001KT) each diluted to 1:500 were used to reveal the C1 and C2 or C3 probes, respectively. The slides were mounted using ProLong^TM^ Gold antifade reagent (Invitrogen). These experiments were imaged with a Spinning Disk Zeiss controlled by Zen Blue software (Olympus, Tokyo, Japan) and processed on the Fiji software (ImageJ) to combine all the images into one containing the maximum intensity information for each pixel. The quantitative analysis was performed on two adjacent sections for at least n=6 per genotype, the results show the mean of the two sections and represent the ratio between the number of dots (representing mRNA strands) and the number of cells (DAPI). For the quantitative analysis along the suture, the analyses encompassed all cells within the suture for *fgfr3^+/+^* fish. In contrast for the *fgfr3^lof/lof^* fish only cells along the bones within the suture were counted, corresponding to the location of the *runx2:*GFP*, sp7:*mCherry*, bglap:*GFP positive cells labelling. The quantitative analysis along the bone surface correspond to the ratio between the number of dots (mRNA strands) and the number of cells (DAPI) bordering above and below the frontal bones.

### Polarization-resolved Second Harmonic Generation microscopy

Multiphoton images of unstained FFPE sections of *fgfr3^+/+^* (n=6) and *fgfr3^lof/lof^* (n=6) fish at the 12SL and 26 SL stages were acquired using a custom-built upright laser scanning setup, as previously described ^52^. Laser excitation was tuned to 860 nm and excitation power was 2 to 8mW at the sample using a 25x, NA 1.05 water immersion objective (XLPLN-MP, Olympus). Both two-photons excited fluorescence (2PEF) and second harmonic generation (SHG) signals were detected in parallel in two trans-detection channels. Endogenous 2PEF enabled to visualize the tissue morphology, while SHG was obtained specifically from collagen fibrils. Polarization-resolved SHG imaging (P-SHG) was performed by recording series of images excited by linear polarizations rotated from 0° to 170° in 10° steps ^52^. Z-stacks of such P-SHG images were recorded at 200 kHz with an axial step of 0.5µm and a pixel size of 420nm. Automated processing by custom written Matlab code then provided: (i) the 2PEF and SHG images averaged over all excitation polarizations, which mitigates the polarization sensitivity; (ii) a pixel-scale orientation map of collagen fibre orientation within the imaging plane ^52^. The latter map was obtained after applying a 2x2 binning filter to increase the signal-to-noise ratio. Pixels with a coefficient of determination less than 0.5 were filtered out ^52^.

Two quantitative parameters were then calculated from these data, keeping only the image from each z-stack that have the highest mean SHG signal. First, the collagen density within the frontal bone was estimated by calculating the mean intensity of the SHG image normalized to the square of the excitation power. Second, the orientation order of the collagen fibres was quantified by computing the circular standard deviation of the orientation distribution obtained from P-SHG images. This disorder metric assesses the width of the orientation distribution using circular statistics; it increases when the orientation disorder increases. To account for the variation of overall orientation of the bone that would bias a measure of absolute local orientation, this circular standard deviation was computed using the P-SHG orientation relative to the orientation of the bone structure, by computing the direction of the frontal bone locally and eliminating small outgrowths.

### Transmission Electron Microscopy

Calvaria from zebrafish at 9, 12 and 20 SL were dissected inside the fixation buffer (for 50mL: 10mL of cacodylate 0.5M, 75µL of glutaraldehyde, 12.5mL of PFA 4% qsp 50mL H2O). The calvarias were left at least 5 days inside the fixation buffer at 4°C and washed 3 times 15 minutes in PBS, then decalcified 24 hrs in EDTA 0.5M pH8 at 4°C. Samples were post fixed in 1% osmium tetroxide 0.1M (Electron Microscopy Science, UK) in 0.1 M Phosphate Buffer (PB) (pH 7.4). Samples were washed 3 times in H2O then dehydrated in alcohol grades: 70% Ethanol 10 min, 90% Ethanol 10 min, 100% Ethanol 3x15min, 100% Propylene oxide (Electron Microscopy Science, UK) 2x5 min. Resin infiltration was performed as following: mix 1:1 Epikote 812: propylene oxide 30 min followed by mix 1:2 Epikote 812: propylene oxide overnight room temperature. Samples were washed in 100% Epikote 812 then embedded in silicon molds in 100% Epikote 812, and Polymerised in a 60°C oven for 24 hours. Ultrathin sections were cut at 90 nm with a Leica UFC7 ultramicrotome (Leica Microsystems GmbH, Germany) and deposed on Gilder grids 200 mesh (Electron Microscopy Science, UK). They were counterstained with uranyl acetate 7% (LFG, France) and Reynold’s lead citrate (LFG, France). Samples were examined in a JEOL 1011 transmission electron microscope (JEOL, Japan) with an ORIUS 1000 CCD camera (GATAN, France), operated with Digital Micrograph software (GATAN, France) for acquisition.

### Statistical analysis

All statistical analyses were performed using GraphPad Prism (version 10, GraphPad Software, La Jolla, CA, USA). Data normality and lognormality were assessed using the D’Agostino & Pearson, Anderson-Darling, Shapiro-Wilk, and Kolmogorov-Smirnov tests. For data following a normal distribution, statistical comparisons were performed using an unpaired t-test when analysing an independent variable between two groups. A two-way ANOVA was used to assess statistically significant differences between three or more independent groups split across two variables. Non-normally distributed data were analysed using a non-parametric Mann-Whitney test for two-group comparisons. A significance threshold of p < 0.05 was applied to all statistical tests. p-values reported in figure legends were considered statistically significant as follows: *, p < 0.05; **, p < 0.01; ***, p < 0.001; and ****, p < 0.0001. All values are presented as mean ± SD.

## Acknowledgments

We are grateful to Dr Chrissy Hammond for providing the Tg(*col2*:mCherry) fish line, Dr Shahad Albadri for the Tg(*7xTCF-Xla.Siam*:GFP), Dr Gilbert Weidinger for providing the Tg(*sp7*:mCherry) and Tg(*bglap*:GFP) fish lines and the members of the fish-facility in the Institut Imagine. We thank all the members of the Institut Imagine Cell Imaging and animal facility. We are grateful to the histology core facility of the Structure Fédérative de Recherche Necker. We thank Paolo Bonaldo for generously providing the antibody against Col VI. We thank Drs. Chris Gordon, Marion Coolen and Sophie Thomas for their valuable advice in writing this manuscript. With regard to funding, this work was supported by ANR-21-CE14-0010 “STARFISH” from the Agence Nationale de le Recherche. We thank the Philanthropy Department of Mutuelles AXA through the “Head and Heart Chair” for supporting the Imagine Institute’s imaging facility. MCSK, AC and YL acknowledge funding from the “Agence Nationale de la Recherche” (ANR-10-INBS-04, ANR-11-EQPX-0029). We thank the ARC Foundation for Cancer Research for the acquisition of the nanozoomer at the histology core facility.

## Authors’ roles

RP designed the experiments, contributed to the acquisition of data, analysed the data and wrote the paper. YM contributed to the acquisition of data. YL and CA contributed to analysis of SHG data. SA and MM contributed to the acquisition of TEM data. MCSK conceived the SHG study, contributed to analysis SHG data and preparation of the paper. FR contributed to analysis of ECM data and preparation of the paper. LLM contributed to analysis of data and preparation of the paper. ED conceived the study, contributed to the analysis of the data and wrote the paper. All authors reviewed the paper and provided intellectual content.

## Supplementary data

**Figure 1 supplement 1.** Fgfr3 is required for coronal suture formation. (A) HE stained sagittal sections of the zebrafish head at the coronal suture level at 20 SL demonstrated that the absence of Fgfr3 prevents bones overlap and coronal suture formation. The dotted lines represent the bone boundaries. Scale bar = 50 µm. (B) Measurement of bone thickness near the coronal suture showed that the *fgfr3^lof/lof^* bones are thicker than *fgfr3^+/+^* bones at the adult stages 20 SL (*fgfr3^+/+^* n=6; *fgfr3^lof/lof^* n=6) and 26 SL (*fgfr3^+/+^* n=7; *fgfr3^lof/lof^* n=6). F: frontal bone. P: Parietal bone. Data are presented as mean ± SD. The p-value were determined by Student’s t tests: ns: not significant; p < 0.05 = *; p < 0.01 = **; p < 0.001 = ***; p < 0.0001 = ****.

**Figure 4 supplement 1.** TRAP-stained sagittal sections of the zebrafish head revealed that TRAP positive cells are present on the viscerocranium (A) and the absence of TRAP positive cells within the metopic suture (B) in both *fgfr3^lof/lof^* and *fgfr3*^+/+^ zebrafish at 12, 20 and 26 SL. The dotted lines represent the bone boundaries. F: Frontal bone. Scale bar = 50 µm.

**Figure 4 supplement 2.** Fgfr3 is an inhibitor of osteoprogenitors expansion within the suture. The nuclei of cells along the suture in *fgfr3^lof/lof^* fish exhibit the shape of osteoprogenitor nuclei. (A) Schematic representation of *fgfr3^+/+^* and *fgfr3^lof/lof^* metopic sutures indicating the different zones studied. (B) High magnification of nuclei stained with DAPI at the osteogenic front (OF) and along the metopic suture of 20 SL *Tg(runx2:GFP*; *fgfr3^+/+ or lof/lof^*) fish. Scale bar = 5 µm. (C) Coronal sections of 20 SL fish imaged by transmission electron microscopy. The dotted lines represent the nuclear boundaries. Scale bar = 2 µm. (D) Quantitative comparison between the nucleus of *fgfr3^+/+^* cells along the suture and the OF and of *fgfr3^lof/lof^* cells along the sutures (*fgfr3^+/+^* n=6; *fgfr3^lof/lof^* n=6). The nuclear shape of *fgfr3^lof/lof^* cells along the suture is similar to that of OF cells in the controls, and not to that of cells at the edge of the metopic suture in the controls. Data are presented as mean ± SD. The p-values were determined by Student’s t tests. ns: not significant; p < 0.05 = *; p < 0.01 = **.

**Figure 4 supplement 3.** (A) Immunofluorescence against GFP (green) and mCherry (red) was performed on metopic sutures of *Tg(sp7:mCherry; bglap:GFP*; *fgfr3^+/+ or lof/lof^*) fish at 26 SL. Nuclei were counterstained with DAPI (blue) and the dotted lines represent the bone boundaries. Scale bar = 50 µm. (B-C) Quantification at 26 SL of the percentage of *sp7*:mCherry positive cells and *bglap*:GFP positive cells along the suture and along the bone surface (*fgfr3^+/+^* n=9; *fgfr3^lof/lof^* n=5). (D) Immunofluorescence against GFP (green) and mCherry (red) was performed on coronal sutures of *Tg(sp7:mCherry; bglap:GFP*; *fgfr3^+/+ or lof/lof^*) fish at 26SL. Nuclei were counterstained with DAPI (blue) and the dotted lines represent the bone boundaries. Scale bar = 50 µm. (E-F) Quantification at 26 SL of the percentage of *sp7*:mCherry and *bglap*:GFP positive cells along the coronal suture (*fgfr3^+/+^* n=5; *fgfr3^lof/lof^* n=5; except *bglap* along the bone surface *fgfr3^lof/lof^* n=4). F: Frontal bone. P: Parietal bone. The p-values were determined by two-way ANOVA. ns: not significant; p < 0.01 = **; p < 0.001 = ***; p < 0.0001 = ****.

**Figure 5 supplement 1.** (A) Expression of *tnmd* and *scx* (red) in coronal sections of the metopic suture at 20 SL, labeled by RNAscope *in situ* hybridization. Nuclei are counterstained with DAPI (blue). Scale bar = 50 µm. (B) Quantitative RNAscope analysis revealed that *tnmd* and *scx* are not differentially expressed in *fgfr3^lof/lof^* compared to controls (*tnmd fgfr3^+/+^* n=8; *fgfr3^lof/lof^* n=6; *scx fgfr3^+/+^* n=8; *fgfr3^lof/lof^* n=7). Data are presented as mean ± SD. F: Frontal bone. p-values were determined by Student’s t tests. ns: not significant. The dotted lines represent the bone boundaries.

**Figure 6 supplement 1.** RNAscope quantitative analyses of *fgfr1a, fgfr1b* and *fgfr2* expression in suture versus bone surface in *fgfr3^+/+^* or *fgfr3^lof/lof^* at 12 and 20 SL showed that *fgfr1a is* more highly expressed along the bone surface than within the suture in *fgfr3^lof/lof^* fish at 12 SL, while *fgfr2* is more highly expressed in bone surface than the suture at 20 SL, in both genotypes (*fgfr1a* and *fgfr1b*: 12 SL: *fgfr3^+/+^* n=6; *fgfr3^lof/lof^* n=6; 20 SL: *fgfr3^+/+^* n=7; *fgfr3^lof/lof^* n=6) (*fgfr2*: 12 SL: *fgfr3^+/+^* n=6; *fgfr3^lof/lof^* n=6; 20 SL: *fgfr3^+/+^* n=7; *fgfr3^lof/lof^* n=7). Data are presented as mean ± SD. The p-values were determined by two-way ANOVA. ns: not significant; p < 0.01 = **; p < 0.001 = ***; p < 0.0001 = ****. The dotted lines represent the bone boundaries.

## References

1. Twigg, S. R. F. & Wilkie, A. O. M. A Genetic-Pathophysiological Framework for Craniosynostosis. Am J Hum Genet 97, 359–377 (2015).

2. Cornille, M., et al. Animal models of craniosynostosis. Neurochirurgie 65, 202–209 (2019).

3. Lana-Elola, E., Rice, R., Grigoriadis, A. E. & Rice, D. P. C. Cell fate specification during calvarial bone and suture development. Developmental Biology 311, 335–346 (2007).

4. Holmes, G., et al. Integrated Transcriptome and Network Analysis Reveals Spatiotemporal Dynamics of Calvarial Suturogenesis. Cell Reports 32, 107871 (2020).

5. Farmer, D. T., et al. Cellular transitions during cranial suture establishment in zebrafish. Nat Commun 15, 6948 (2024).

6. Maruyama, T., Jeong, J., Sheu, T.-J. & Hsu, W. Stem cells of the suture mesenchyme in craniofacial bone development, repair and regeneration. Nat Commun 7, 10526 (2016).

7. Zhao, H., et al. The suture provides a niche for mesenchymal stem cells of craniofacial bones. Nat Cell Biol 17, 386–396 (2015).

8. Maruyama, T., et al. BMPR1A maintains skeletal stem cell properties in craniofacial development and craniosynostosis. Sci. Transl. Med. 13, eabb4416 (2021).

9. Wilk, K., et al. Postnatal Calvarial Skeletal Stem Cells Expressing PRX1 Reside Exclusively in the Calvarial Sutures and Are Required for Bone Regeneration. Stem Cell Reports 8, 933–946 (2017).

10. Debnath, S., et al. Discovery of a periosteal stem cell mediating intramembranous bone formation. Nature 562, 133–139 (2018).

11. Teng, C. S., et al. Altered bone growth dynamics prefigure craniosynostosis in a zebrafish model of Saethre-Chotzen syndrome. Elife 7, e37024 (2018).

12. Bok, S., et al. A multi-stem cell basis for craniosynostosis and calvarial mineralization. Nature 621, 804–812 (2023).

13. Herring, S. W. Mechanical influences on suture development and patency. Front Oral Biol 12, 41– 56 (2008).

14. Izu, Y., Ezura, Y., Koch, M., Birk, D. E. & Noda, M. Collagens VI and XII form complexes mediating osteoblast interactions during osteogenesis. Cell Tissue Res 364, 623–635 (2016).

15. Izu, Y., et al. Type XII collagen regulates osteoblast polarity and communication during bone formation. J Cell Biol 193, 1115–1130 (2011).

16. Huang, X., et al. Gli1+ Cells Residing in Bone Sutures Respond to Mechanical Force via IP3R to Mediate Osteogenesis. Stem Cells Int 2021, 8138374 (2021).

17. Jing, D., et al. Response of Gli1+ Suture Stem Cells to Mechanical Force Upon Suture Expansion. J Bone Miner Res 37, 1307–1320 (2022).

18. Daponte, V., et al. Cell differentiation and matrix organization are differentially affected during bone formation in osteogenesis imperfecta zebrafish models with different genetic defects impacting collagen type I structure. Matrix Biol 121, 105–126 (2023).

19. Ga, M., Sj, O., Ir, M., Ka, P. & Ew, J. Fibroblast growth factor receptor 3 (FGFR3) transmembrane mutation in Crouzon syndrome with acanthosis nigricans. Nature genetics 11, (1995).

20. Muenke, M., et al. A unique point mutation in the fibroblast growth factor receptor 3 gene (FGFR3) defines a new craniosynostosis syndrome. Am J Hum Genet 60, 555–564 (1997).

21. Makrythanasis, P., et al. A novel homozygous mutation in FGFR3 causes tall stature, severe lateral tibial deviation, scoliosis, hearing impairment, camptodactyly, and arachnodactyly. Hum. Mutat. 35, 959–963 (2014).

22. Deng, C., Wynshaw-Boris, A., Zhou, F., Kuo, A. & Leder, P. Fibroblast growth factor receptor 3 is a negative regulator of bone growth. Cell 84, 911–921 (1996).

23. Nah, H.-D., Koyama, E., Agochukwu, N. B., Bartlett, S. P. & Muenke, M. Phenotype profile of a genetic mouse model for Muenke syndrome. Childs Nerv Syst 28, 1483–1493 (2012).

24. Cornille, M., et al. FGFR3 overactivation in the brain is responsible for memory impairments in Crouzon syndrome mouse model. J Exp Med 219, e20201879 (2022).

25. Laue, K., et al. Craniosynostosis and multiple skeletal anomalies in humans and zebrafish result from a defect in the localized degradation of retinoic acid. Am. J. Hum. Genet. 89, 595–606 (2011).

26. Pereur, R. & Dambroise, E. Insights into Craniofacial Development and Anomalies: Exploring Fgf Signaling in Zebrafish Models. Curr Osteoporos Rep 22, 340–352 (2024).

27. Sun, X., et al. Fgfr3 mutation disrupts chondrogenesis and bone ossification in zebrafish model mimicking CATSHL syndrome partially via enhanced Wnt/β-catenin signaling. Theranostics 10, 7111–7130 (2020).

28. Dambroise, E., et al. Fgfr3 Is a Positive Regulator of Osteoblast Expansion and Differentiation During Zebrafish Skull Vault Development. J Bone Miner Res 35, 1782–1797 (2020).

29. Izu, Y. & Birk, D. E. Collagen XII mediated cellular and extracellular mechanisms in development, regeneration, and disease. Front. Cell Dev. Biol. 11, 1129000 (2023).

30. Miao, K. Z., Cozzone, A., Caetano-Lopes, J., Harris, M. P. & Fisher, S. Osteoclast activity sculpts craniofacial form to permit sensorineural patterning in the zebrafish skull. Front. Endocrinol. 13, 969481 (2022).

31. Farmer, D. T., et al. The developing mouse coronal suture at single-cell resolution. Nat Commun 12, 4797 (2021).

32. Moshkovsky, A. R. & Kirschner, M. W. The nonredundant nature of the Axin2 regulatory network in the canonical Wnt signaling pathway. Proc. Natl. Acad. Sci. U.S.A. 119, e2108408119 (2022).

33. Bobzin, L., et al. FGFR2 directs inhibition of WNT signaling to regulate anterior fontanelle closure during skull development. Development 152, dev204264 (2025).

34. Di Rocco, F., et al. FGFR3 mutation causes abnormal membranous ossification in achondroplasia. Hum. Mol. Genet. 23, 2914–2925 (2014).

35. Kague, E., et al. Osterix/Sp7 limits cranial bone initiation sites and is required for formation of sutures. Dev. Biol. 413, 160–172 (2016).

36. Katsianou, M. A., Adamopoulos, C., Vastardis, H. & Basdra, E. K. Signaling mechanisms implicated in cranial sutures pathophysiology: Craniosynostosis. BBA Clinical 6, 165–176 (2016).

37. Jiang, T., Ge, S., Shim, Y. H., Zhang, C. & Cao, D. Bone morphogenetic protein is required for fibroblast growth factor 2-dependent later-stage osteoblastic differentiation in cranial suture cells. Int J Clin Exp Pathol 8, 2946–2954 (2015).

38. Warren, S. M., Brunet, L. J., Harland, R. M., Economides, A. N. & Longaker, M. T. The BMP antagonist noggin regulates cranial suture fusion. Nature 422, 625–629 (2003).

39. Matsushita, T., et al. FGFR3 promotes synchondrosis closure and fusion of ossification centers through the MAPK pathway. Hum. Mol. Genet. 18, 227–240 (2009).

40. Ng, J. Q., et al. Loss of Grem1-lineage chondrogenic progenitor cells causes osteoarthritis. Nat Commun 14, 6909 (2023).

41. Ohbayashi, N., et al. FGF18 is required for normal cell proliferation and differentiation during osteogenesis and chondrogenesis. Genes Dev. 16, 870–879 (2002).

42. Menon, S., et al. Skeletal stem and progenitor cells maintain cranial suture patency and prevent craniosynostosis. Nat Commun 12, 4640 (2021).

43. Mirando, A. J., Maruyama, T., Fu, J., Yu, H.-M. I. & Hsu, W. β-catenin/cyclin D1 mediated development of suture mesenchyme in calvarial morphogenesis. BMC Dev Biol 10, 116 (2010).

44. Reinhold, M. I. & Naski, M. C. Direct interactions of Runx2 and canonical Wnt signaling induce FGF18. J Biol Chem 282, 3653–3663 (2007).

45. Gregory, C. A., Ma, J. & Lomeli, S. The coordinated activities of collagen VI and XII in maintenance of tissue structure, function and repair: evidence for a physical interaction. Front. Mol. Biosci. 11, 1376091 (2024).

46. Izu, Y., et al. Collagen XII mediated cellular and extracellular mechanisms regulate establishment of tendon structure and function. Matrix Biology 95, 52–67 (2021).

47. Parichy, D. M., Elizondo, M. R., Mills, M. G., Gordon, T. N. & Engeszer, R. E. Normal table of postembryonic zebrafish development: staging by externally visible anatomy of the living fish. Dev. Dyn. 238, 2975–3015 (2009).

48. Knopf, F., et al. Bone regenerates via dedifferentiation of osteoblasts in the zebrafish fin. Dev. Cell 20, 713–724 (2011).

49. Moro, E., et al. In vivo Wnt signaling tracing through a transgenic biosensor fish reveals novel activity domains. Dev Biol 366, 327–340 (2012).

50. Tonelotto, V., et al. Collagen VI ablation in zebrafish causes neuromuscular defects during developmental and adult stages. Matrix Biol 112, 39–61 (2022).

51. Bader, H. L., et al. Zebrafish collagen XII is present in embryonic connective tissue sheaths (fascia) and basement membranes. Matrix Biol 28, 32–43 (2009).

52. Raoux, C., et al. Quantitative structural imaging of keratoconic corneas using polarization-resolved SHG microscopy. Biomed Opt Express 12, 4163–4178 (2021).

