## Supplementary figures and images for "Cranial suture integrity is maintained by Fgfr3 in zebrafish"

### Figure 1 supplement 1

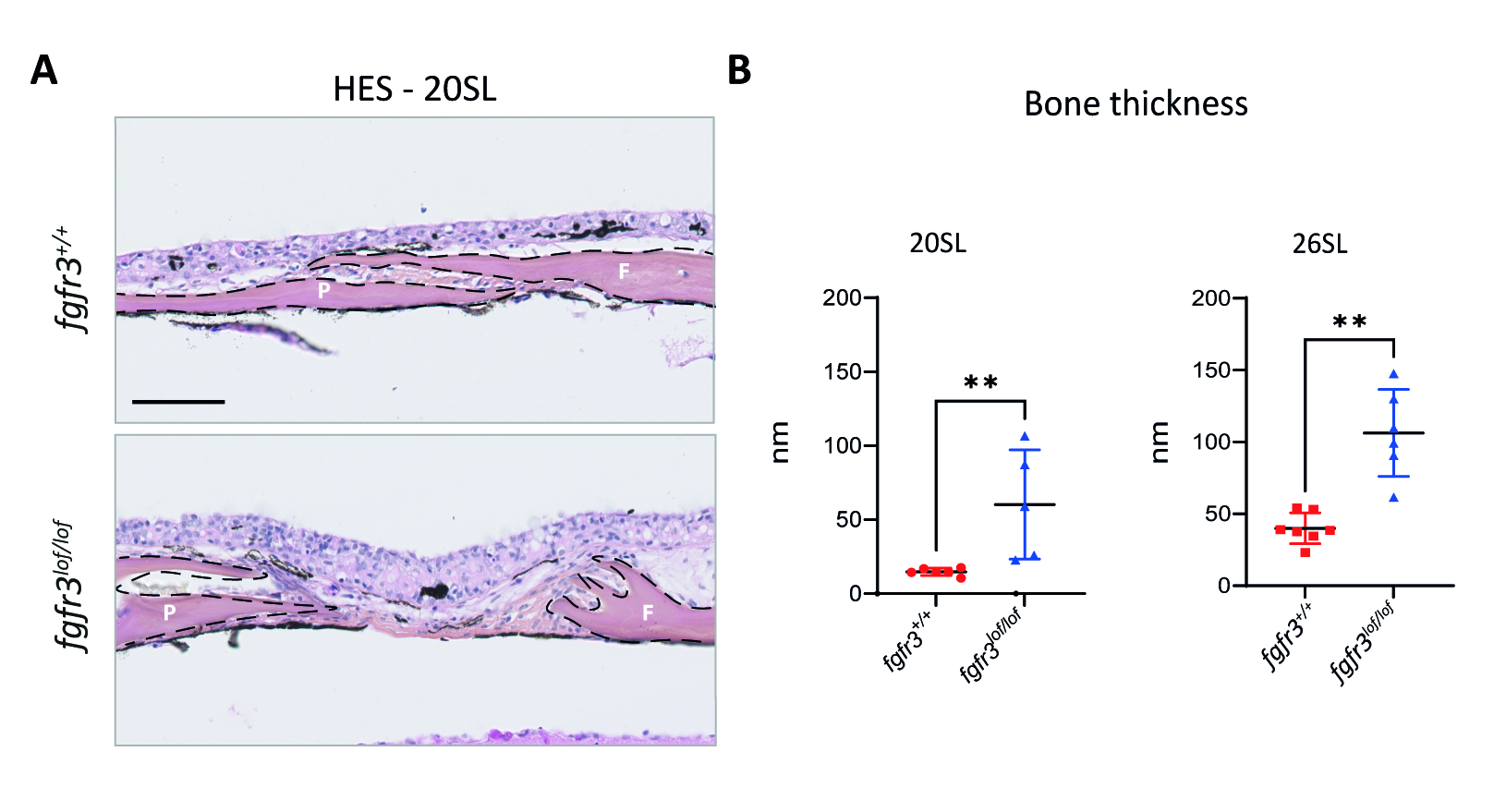

### Figure 4 supplement 1

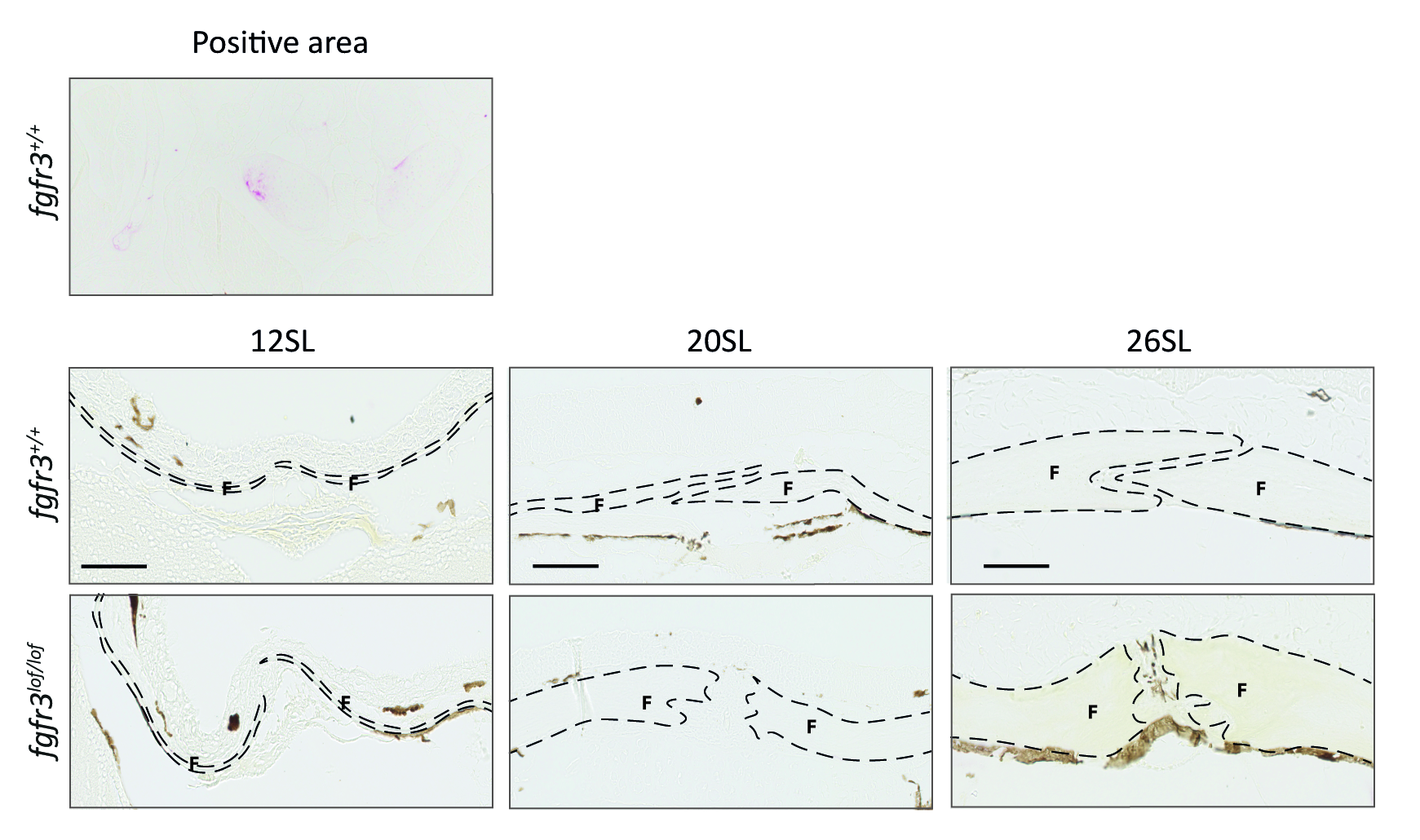

### Figure 4 supplement 2

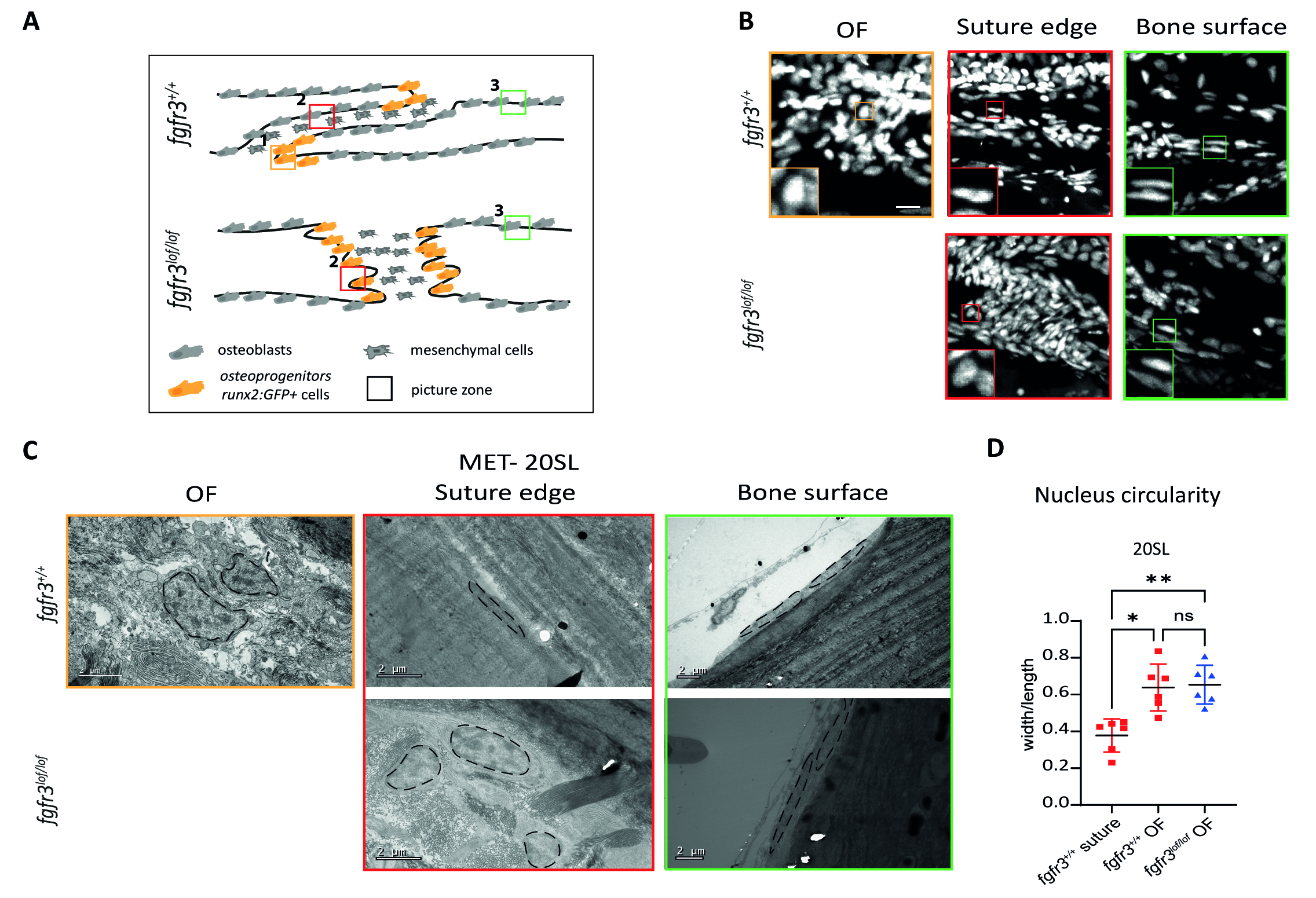

### Figure 4 supplement 3

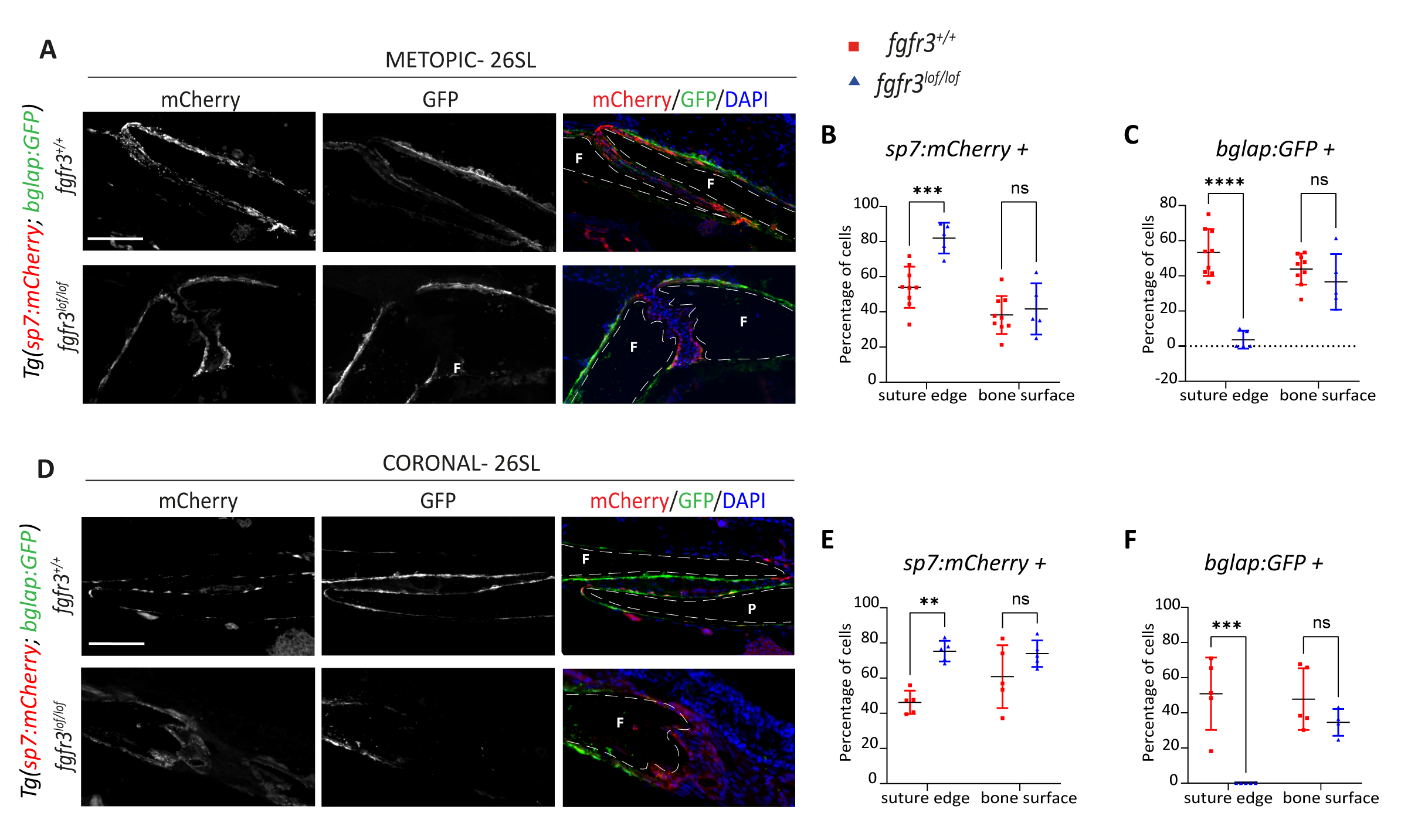

### Figure 5 supplement 1

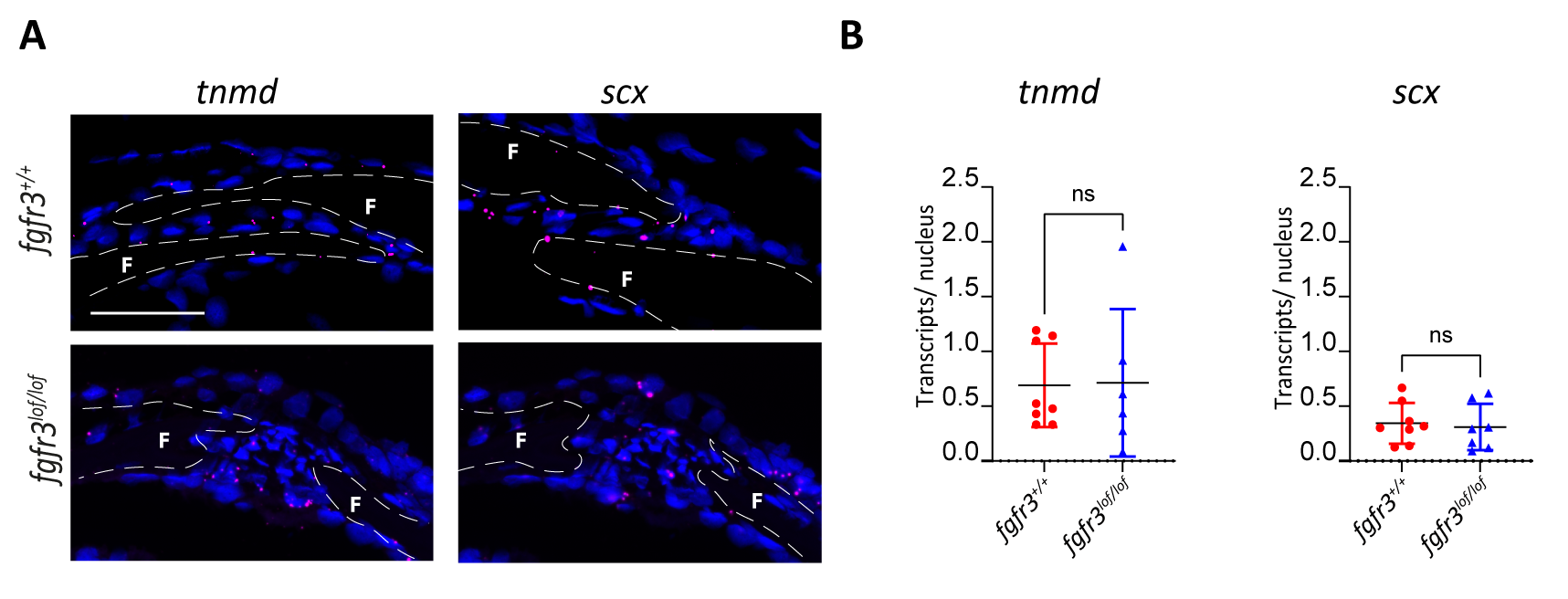

### Figure 6 supplement 1

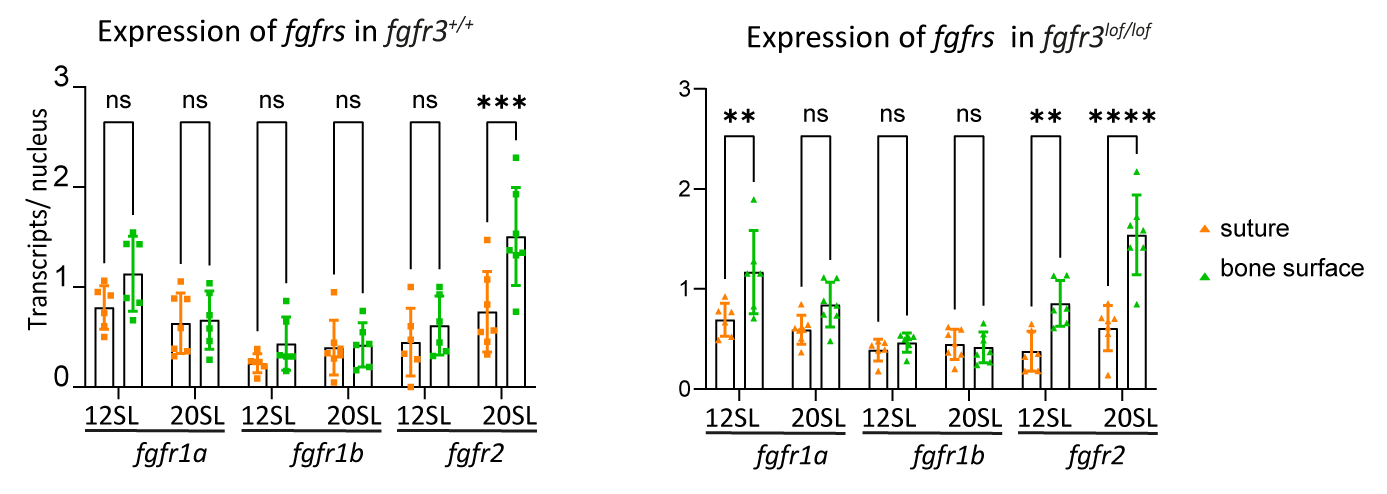
